# Novel laminarin-binding CBMs in multimodular proteins of marine *Bacteroidota* feature prominently in phytoplankton blooms

**DOI:** 10.1101/2023.09.07.556657

**Authors:** Marie-Katherin Zühlke, Elizabeth Ficko-Blean, Daniel Bartosik, Nicolas Terrapon, Alexandra Jeudy, Murielle Jam, Fengqing Wang, Norma Welsch, Robert Larocque, Diane Jouanneau, Tom Eisenack, François Thomas, Anke Trautwein-Schult, Hanno Teeling, Dörte Becher, Thomas Schweder, Mirjam Czjzek

**Author notes:** corresponding authors: Mirjam Czjzek, Marie-Katherin Zühlke.

## Abstract

The ß-(1,3)-glucan laminarin functions as storage polysaccharide in marine stramenophiles such as diatoms. Laminarin is abundant, water-soluble and structured simply, making it an attractive substrate for marine bacteria. As a consequence, many marine bacteria have developed competitive strategies to scavenge and decompose laminarin, which involves carbohydrate-binding modules (CBMs) as key players. We therefore functionally and structurally characterized two yet unassigned domains as laminarin-binding CBMs in multimodular proteins from our model bacterium *Christiangramia forsetii* KT0803^T^, hereby unveiling the novel laminarin-binding CBM families CBMxx and CBMyy (official CAZy numbering will be provided upon acceptance of the manuscript in a peer-reviewed journal). We discovered four CBMxx repeats in a surface glycan-binding protein (SGBP) and a single CBMyy combined with a glycoside hydrolase module from family 16 (GH16_3). Our analyses revealed that both modular proteins have an elongated shape, and that the GH16_3 displayed a higher flexibility than the SGBP. While motility of both polypeptide chains may facilitate recognition and/or degradation of laminarin, constraints in the SGBP may support docking of laminarin onto the bacterial surface. The exploration of bacterial metagenome-assembled genomes (MAGs) from phytoplankton blooms in the North Sea revealed that both laminarin-binding CBM families are widely distributed among marine *Bacteroidota*, illustrating the high adaptability of modularity in sugar-binding and -degrading proteins. High expression of CBMxx- and CBMyy-containing proteins during phytoplankton blooms further underpins their importance in marine laminarin usage.

## Introduction

Laminarin is an important energy and carbon storage compound in marine stramenophiles, including abundant diatoms. Therefore, huge amounts of laminarin are produced during common diatom-dominated phytoplankton blooms (1). Laminarin is composed of a ß-(1,3)-linked glucose backbone with occasional ß-(1,6)-linked glucose branches that differ in length and frequency (2, 3). Due to its abundance, water solubility and simple structure, marine bacteria often prefer laminarin as a substrate over less soluble and more complex polysaccharides (4, 5). Polysaccharide utilization loci (PULs) dedicated to laminarin breakdown are thus both widely distributed (6, 7) and are highly expressed (4, 8–10) in marine bacteria that thrive during phytoplankton blooms. PULs are distinct genetic loci that encode proteins to sense, digest and transport sugars. *Bacteroidota* are key players in marine laminarin turnover and many glycoside hydrolases (GHs) active on laminarin from members of this phylum have been characterized, such as GH16 and GH17 family representatives that act on the ß-(1,3)-main chain, or GH30 family representatives that remove ß-(1,6)-side chains (11–14). Some of these GHs are bound to the cell surface, where they degrade laminarin into smaller-sized oligosaccharides suitable for uptake and transport (11). In *Bacteroidota*, the oligosaccharide shuttle to the periplasm is mediated by an accessory sugar-binding SusD-like protein, tethered to the outer membrane, and a SusC-like TonB-dependent transporter (TBDT) (reviewed in 15, 16). In complex with their target substrate, SusD-like proteins act as a hinge to relay their cargo into the SusC-like TBDT (17). In addition to SusD-like proteins, bacteria may have additional surface glycan-binding proteins (SGBPs), which are often encoded directly downstream of the *susCD*-like gene tandem and are sometimes termed ‘SusE-positioned’ proteins. The designations ‘-like’ *versus* ‘-positioned’ intend to distinguish between homologous *versus* analogous proteins, and refer to the starch utilization system (Sus) (18, 19), which marked the beginning of the era of PUL exploration. While SusD-like proteins have a single binding site and are almost entirely α-helical, SusE-positioned SGBPs may have several carbohydrate-binding modules (CBMs), which often display a ß-sandwich fold structure and function as carbohydrate docking sites. Since SusD-like proteins also classify as SGBPs, they have sometimes been named ‘SGBP-A’ in the literature, whereas CBM-containing SGBPs are named ‘SGBP-B’ (20). Some examples of mixed-linkage ß-glucan and laminarin-related SusD-like proteins and CBM-containing SGBPs have recently been thoroughly analyzed in gut strains from the *Bacteroides* genus (21). By comparison, there is only one structure for a marine laminarin-binding SusD-like protein, which recognizes ß-(1,6)-branches in laminarin (22), and marine examples of CBM-containing SGBPs that bind laminarin are lacking. As aforementioned, unlike SusD-like proteins, CBM-containing SGBPs rarely show detectable similarity/homology (20, 21) and are thus difficult to predict. Therefore, CBM-containing SGBPs are largely overlooked and probably underestimated. In addition to SusD-like proteins and CBM-containing SGBPs, laminarin-binding CBMs are often linked to enzymes within multidomain structures (23, 24), which further underlines the importance of CBMs in marine laminarin degradation.

The laminarin-consuming *Christiangramia forsetii* KT0803^T^ (formerly *Gramella forsetii* (25)), a member of the *Bacteroidota*, has been isolated during a phytoplankton bloom in the North Sea (26). A GH16_3 and a ‘conserved hypothetical protein’, encoded in the *C. forsetii* laminarin PUL, were among the most abundantly produced proteins when growing on laminarin as a sole carbon source (27). Based on subproteome analyses it was suggested that the two proteins are tethered to the outer membrane and it was speculated that the ‘conserved hypothetical protein’, which is SusE-positioned, might function as an SGBP (27).

This study aimed to illuminate the relevance of laminarin-binding CBMs in the scavenging and processing of laminarin, which is abundantly produced during phytoplankton blooms. We therefore structurally and functionally characterized an SGBP and a GH16_3 from the marine model bacterium *C. forsetii*, both of which display novel laminarin-binding CBMs in their multimodular structure. This gave insights into specific adaptations of a non-catalytic and a catalytic protein, respectively, with respect to their roles in first steps of laminarin utilization. It further allowed us to illustrate the distribution of these CBMs in marine bacteria and their expression as a response to algae proliferation.

## Results

### Multidomain proteins Cf-SGBP and Cf-GH16_3

We investigated two proteins that are encoded in a laminarin PUL and that were highly produced in laminarin-grown cells of *C. forsetii* (27): a putative CBM-containing SGBP (Cf-SGBP, locus tag GFO_RS17395) and a protein containing a GH16_3 catalytic module (Cf-GH16_3, locus tag GFO_RS16360) (Fig. 1a, further information on domain nomenclature is provided in the Materials and Methods or Fig. 1b). Subproteome analyses (27) and the presence of a lipoprotein signal peptide support that both proteins are tethered to the outer membrane. Structural prediction using Alphafold2 (AF) (28) and Phyre2 (29) uncovered that the Cf-SGBP displays two Ig-like folded (ß-sandwich) modules at its N-terminus followed by four putative CBMs (I-IV) (Fig. 1b) with high sequence-similarity (47-84%), suggesting internal duplication events. The Cf-GH16_3 on the other hand is predicted to feature a single N-terminal Ig-like folded module followed by a putative CBM, with no significant similarity to CBMs of the Cf-SGBP, and the C-terminal GH16_3 catalytic domain at the C-terminus.

**Fig. 1.**
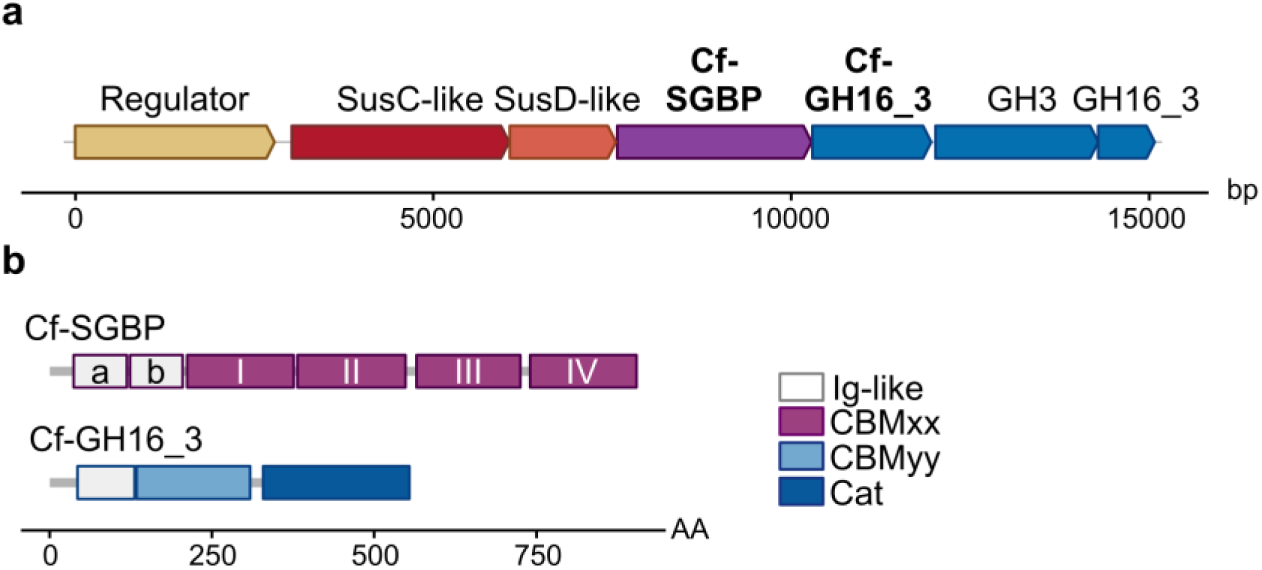
The laminarin PUL of *Christiangramia forsetii* encodes the multimodular Cf-SGBP and Cf-GH16_3 containing the novel CBMxx and CBMyy, respectively. **a** The *C. forsetii* laminarin PUL encodes multidomain proteins, including a novel CBM-containing SGBP (Cf-SGBP) and a GH16_3 (Cf-GH16_3), highlighted bold. **b** Domain organization of the Cf-SGBP and Cf-GH16_3. Here, gray lines represent N-terminal lipoprotein signal peptides or linkers between modules. The Cf-SGBP displays two Ig-like fold modules (a,b) followed by four CBMxxs (I-IV). The Cf-GH16_3 comprises an Ig-like fold module and CBMyy in addition to its catalytic module (Cat).

All putative CBM sequences were submitted to Dali searches (30), which did not reveal any close relatives with solved 3D structures. The four CBMs from the Cf-SGBP were only remotely similar to a module attached to a GH50 from *Saccharophagus degradans* (PDB ID 4BQ2) (31), with a z-score >12 and 10-15% identity. To date, this module is not classified in any CBM family in the CAZy classification, in the absence of a characterized close homolog. Similarly, the CBM of the Cf-GH16_3 returned uncharacterized proteins, including hypothetical CBMs, as best hits with z-scores ∼8 and 9-12% sequence identity (PDB ID 4L3R and 4QRL).

Sequence analysis confirmed that the putative *C. forsetii* CBMs did not exhibit any domains belonging to a known CBM family. Hence, the functional characterization of the CBMs of the Cf-SGBP and Cf-GH16_3 (cf two following sections below) provided the basis for founding the novel CAZy (32) CBM families CBMxx and CBMyy, respectively (official CAZy numbering will be provided upon acceptance of the manuscript in a peer-reviewed journal). We determined the 3D crystal structure at 2.0 Å resolution for the isolated C-terminal CBMxx, Cf-SGBP-CBMxx_IV_, in complex with laminaritriose by protein crystallography (Fig. 2a, Table S1). Least square superimposition of the corresponding AF predicted structure to the 3D crystal structure using COOT (33) and matching all atoms leads to an RMSD of 1.765. This high agreement (Fig. 2b) supported reliability for the AF predictions for the other CBMxxs, which further indicated two things: the four CBMxxs are highly similar in fold and CBMyy differs from CBMxx. The crystal structure of the Cf-SGBP-CBMxx_IV_ reveals a typical CBM ß-sandwich fold, consisting of two antiparallel ß-sheets composed of four or five ß-strands that form a concave and convex face, respectively (Fig. 2a). The concave face forms a deep and narrow binding cleft, also established by two expansive loops. While the concave face contains residues mediating polar interactions with the substrate, distal loops contain aromatic residues (Fig. 2a) to establish CH-π interactions (34). AF predicted a similar fold for all four CBMxxs (Fig. 2c) and residues that mediate interaction with laminaritriose in the Cf-SGBP-CBMxx_IV_ align (Fig. 2d). Remarkably, differences are seen in Cf-SGBP-CBMxx_III_ loop 2, which results in a wider and very shallow binding cleft (Fig. 2c and d, Fig. S1).

**Fig. 2.**
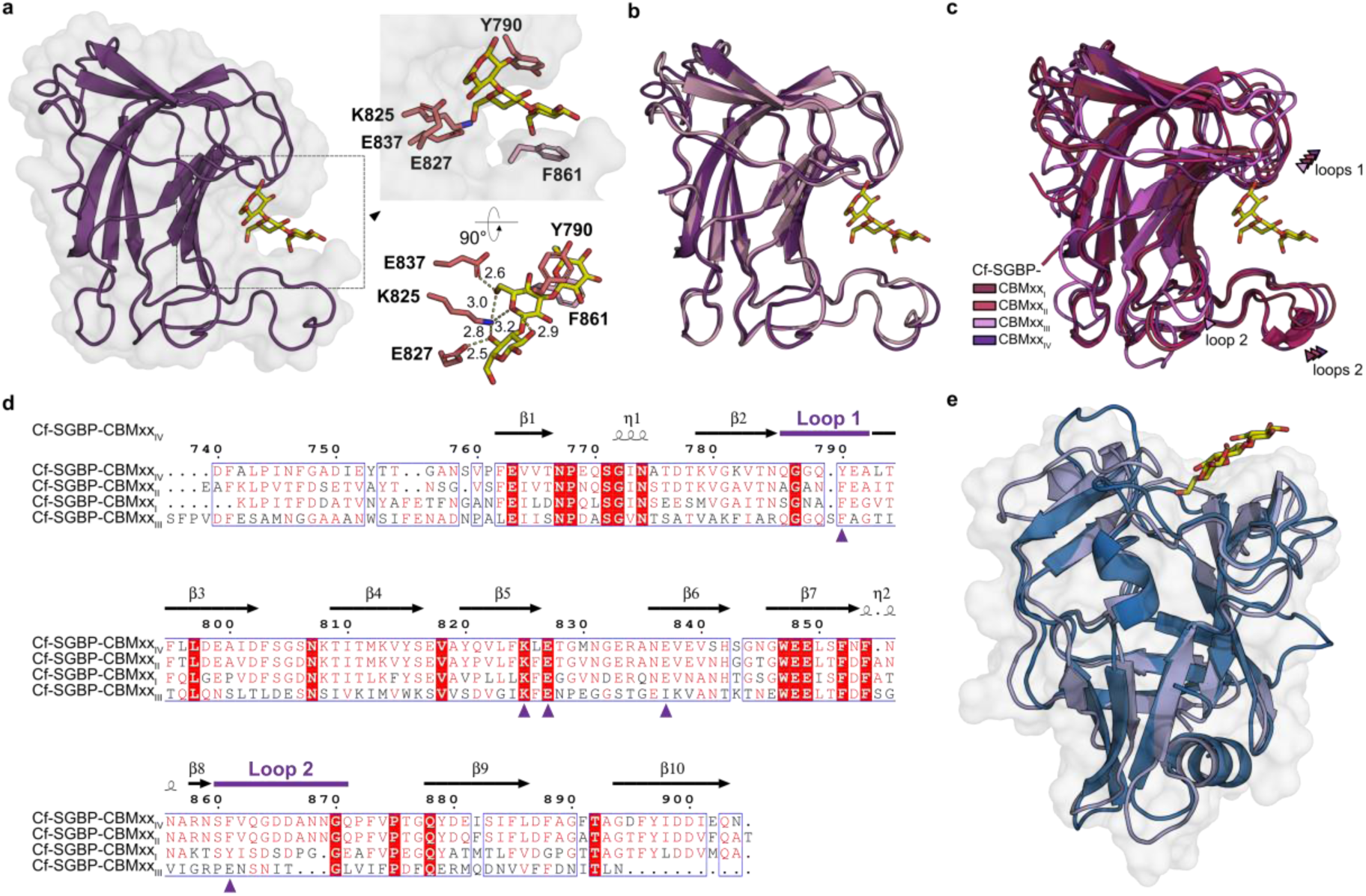
CBMxx repeats of the Cf-SGBP and CBMyy of the Cf-GH16_3: binding clefts *versus* platform. **a** 3D crystal structure of the Cf-SGBP-CBMxx_IV_ with laminaritriose, highlighting interacting residues. **b** Overlay of the crystal structure of the Cf-SGBP-CBMxx_IV_ (violet) and the corresponding AF prediction (light pink), supporting the accuracy of AF. **c** Overlay of AF predicted CBMxx repeats (I-III) with the crystal structure of the Cf-SGBP-CBMxx_IV_. In particular Cf-SGBP-CBMxx_III_ differs in one of the loops delimiting the binding groove, which results in a flat binding region flanked only by one loop (see also Fig. S1). **d** Alignment of the Cf-SGBP-CBMxxs visualized using ESPript 3.0 (https://espript.ibcp.fr) (35). Identical residues are highlighted in red boxes, similar residues using red letters. Residues involved in substrate binding in the Cf-SGBP-CBMxx_IV_ are indicated by violet arrow heads; the two loops flanking the substrate binding groove are indicated by violet lines above the alignment. **e** CBMyy of the Cf-GH16_3 displays a ß-barrel structure, missing a distinct binding cleft compared to CBMxx. The AF predicted model (blue) superimposes very closely to a CBM of an SGBP from *Bacteroidetes fluxus* (21) binding laminarin and mixed-linkage ß(1,3)/ß(1,4)-glucan on a platform (lightblue, here laminaritriose, PDB ID 7KV7, see also Fig. S2).

Compared to the ß-sandwich-fold of CBMxx, CBMyy rather displays a ß-barrel structure with two additional α-helices and an additional β-strand pair. Together with a lacking binding groove, the *C. forsetii* CBMyy is structurally related to a CBM present in a *Bacteroidetes fluxus* SGBP (Fig. 2e, Fig. S2) (21) devoid of a catalytic GH16_3. Our analyses reveal that the *B. fluxus* CBM classifies as a CBMyy.

### CBMxx and CBMyy bind laminarin

Affinity gel electrophoresis (AGE) demonstrated that the Cf-SGBP and Cf-GH16_3 bind laminarin. We narrowed down binding to the CBMs of both proteins, and excluded binding of the N-terminal Ig-like folded modules (Fig. 3a, Fig. S3 and S4). Binding to laminarin induced a distinct shift of the band for the full-length Cf-SGBP (Fig. 3b, lane 6), whereas it induced a small but distinct shift for the full-length Cf-GH16_3 with an altered, ‘mustache’-like band shape (Fig. 3b, lane 1). In comparison, the binding of the isolated catalytic domain Cf-GH16_3-Cat could not be inferred from AGE, nor was an altered band shape detected (Fig. 3b, lane 2). We hypothesize that the catalytic module releases smaller laminarin oligosaccharides that diffuse from the gel, which is why no shift is obtained for the dissected active catalytic module. Accordingly, the shift was restored with mutants of the dissected catalytic domain (Cf-GH16_3-Cat_E442S_ and Cf-GH16_3-Cat_E447S_), but without the detection of the altered band shape (Fig. 3b, lanes 3 and 4). Unexpectedly, we found that the GH16_3-catalytic activity reduced the gel shift of the Cf-SGBP and resulted in a mustache-like band shape of the Cf-SGBP wherever the active enzyme was present (Fig. 3b, lanes 7 and 8). This alteration was not obtained for the Cf-SGBP loaded together with the catalytic mutants (Fig. 3b, lanes 9 and 10). We suggest that in the presence of active enzyme, the CBM-containing polypeptides (Fig. 3b, lanes 1, 7 and 8) bind to residual laminarin in-between the gel lanes, thereby modifying the shape of the bands and resulting in the ‘mustache’ effect.

**Fig. 3.**
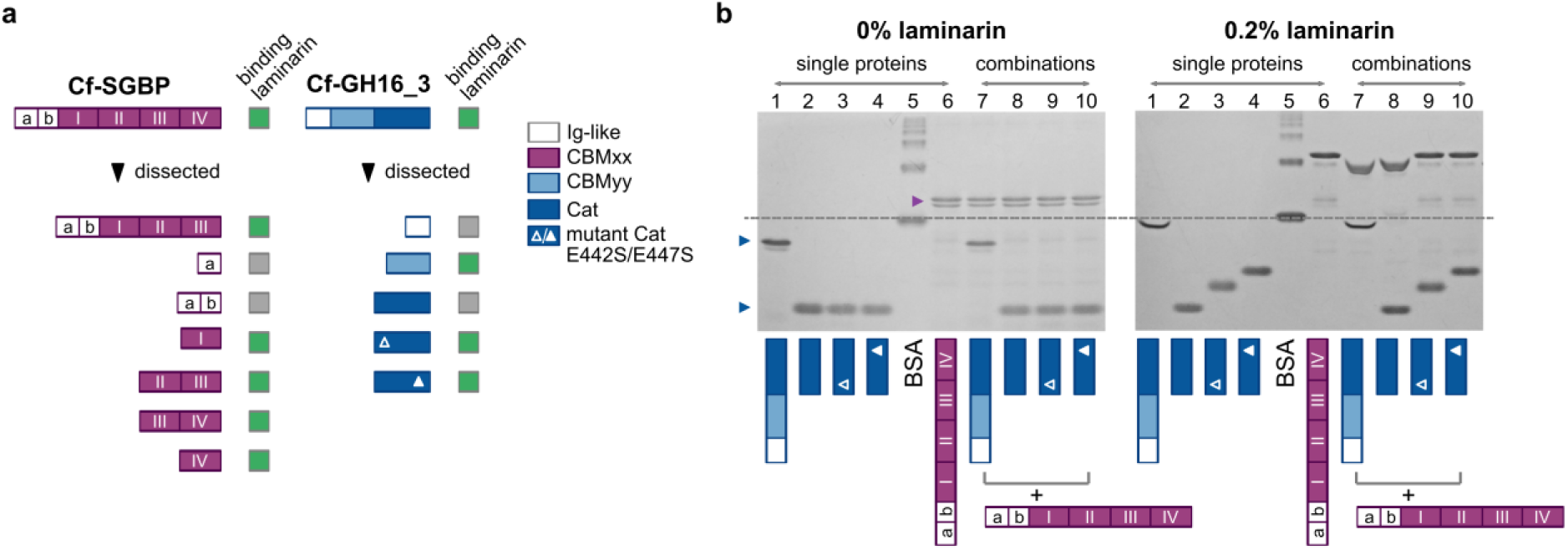
Identification of laminarin-binding domains in the Cf-SGBP and Cf-GH16_3. **a** Summary of AGE runs, where green boxes indicate binding, which was concluded from corresponding gels in Fig. 3b, Fig. S3 and S4. **b** AGE of the Cf-GH16_3 revealed catalytic activity during runs. The Cf-GH16_3, the dissected catalytic module Cf-GH16_3-Cat and two mutants thereof (Cf-GH16_3-Cat_E442S_ and Cf-GH16_3-Cat_E447S_) were loaded individually (lanes 1-4) or in combination with the non-catalytic Cf-SGBP (lanes 7-10), compared to the Cf-SGBP alone (lane 6). While a shift in migration in laminarin gels indicated binding (using BSA as a control, lane 5), an altered ‘mustache’-like band shape of CBM-containing proteins in laminarin gels (lane 1, 7 and 8) revealed the presence of GH16_3 activity within the lane. Detailed information is provided in the main text.

In addition to laminarin binding, AGE also demonstrated binding of the mixed-linkage ß(1,3)/ß(1,4)-glucan (MLG) lichenan by Cf-GH16_3-CBMyy (Fig. S5). Activity of the Cf-GH16_3 on lichenan was not detectable by reducing sugar assay (RSA); however, fluorophore-assisted carbohydrate electrophoresis (FACE) analyses demonstrated weak activity (Fig. S6).

### Multiple CBMxx domains in the Cf-SGBP: is ‘more’ really ‘better’?

While the *C. forsetii* Cf-GH16_3 contains a single CBMyy to bind laminarin, the Cf-SGBP contains four consecutive CBMxxs. AGE did not provide indications on whether the higher number of CBMs in the Cf-SGBP translates into a higher binding efficiency (Fig. S3). In order to explore putative benefits of multiple CBMxx domains in more detail, we investigated laminarin binding by isothermal titration calorimetry (ITC). We selected the full-length Cf-SGBP, the tandems Cf-SGBP-CBMxx_II/III_ and Cf-SGBP-CBMxx_III/IV_, as well as the single module Cf-SGBP-CBMxx_IV_, for which we had a 3D crystal structure, to cover a range of CBMxx repetitions for comparisons. Other constructs were either insufficiently stable or insufficiently produced and thus unfortunately not available for ITC measurements.

ITC confirmed laminarin-binding for the selected proteins (Table 1, Fig. S7a). Unexpectedly, the highest affinity for laminarin was determined for the individual Cf-SGBP-CBMxx_IV_ (1.04 x 10^5^ M^-1^), which exceeded affinity of the other constructs by an order of magnitude (Table 1) and is in the same range as described for the N-terminal CBM6 of a GH16_3 (LamC) in *Zobellia galactanivorans* Dsij^T^ (23). A binding stoichiometry (n) of close to 0.5 indicates two CBMxx_IV_s are able to interact with one ligand molecule. The binding stoichiometry is increased to ∼0.75 for the double module constructs and the full-length Cf-SGBP binds 1.6 laminarin molecules per polypeptide chain, though no significantly increased affinity was seen. Instead, affinities of the full-length Cf-SGBP and the two CBMxx tandems were relatively similar (Cf-SGBP 3.09 x 10^4^ M^-1^, Cf-SGBP-CBMxx_II/III_ 2.76 x 10^4^ M^-1^ and Cf-SGBP-CBMxx_III/IV_ 3.63 x 10^4^ M^-1^). We speculate that the increased affinity of the individual CBMxx_IV_ for laminarin relative to the tandems is provided by the absence of steric/spatial inhibition of binding from the other CBMxx. Thus, in the absence of the other modules, two individual CBMxx_IV_ are more optimally accommodated on the same laminarin chain. Conversely, the steric hindrance of successive CBMs in the Cf-SGBP may contribute to partial binding site inaccessibility. Furthermore, the longer constructs pay more entropic penalties than the single Cf-SGBP-CBMxx_IV_, probably related to the loss of conformational freedom upon ligand binding. Notably, the Cf-SGBP-CBMxx_IV_ also bound laminarin-derived oligosaccharides DP7 with a higher affinity than DP5, although with a lower affinity than laminarin (Table 1, Fig. S7b). This is coherent with increased affinity of the individual Cf-SGBP-CBMxx_IV_ on laminarin, as it suggests that this CBM can accommodate more than DP7 in the ligand binding site. For all interactions, binding was enthalpy-driven, with an entropic penalty (Table 1). Confirming our results from AGE, binding of laminarin by the Cf-SGBP-Ig_a_ was not detected.

**Table 1.** Affinity of laminarin (Lam) and its oligosaccharides (DP7 and DP5) to the Cf-SGBP and/or dissected modules thereof. Mean values and standard deviation (±) of ITC experiments (n≥3). Measurements were performed at 293.15 K. For each run, ΔG was calculated (Gibbs-Helmholtz equation, ΔG=ΔH-TΔS).

| **Cf-SGBP and selected modules - laminarin** |  |  |  |  |  |
| --- | --- | --- | --- | --- | --- |
| | $K_a$<br>[$\text{M}^{-1}$] | $\Delta H$<br>[kcal mol<sup>-1</sup>] | $\Delta S$<br>[cal mol<sup>-1</sup> K<sup>-1</sup>] | N<br>[sites] | $\Delta G$<br>[kcal mol<sup>-1</sup>] |
| Cf-SGBP | $3.09 (\pm 0.14) \times 10^4$ | $-17.3 \pm 0.11$ | $-38.53 \pm 0.46$ | $1.60 \pm 0.05$ | $-6.02 \pm 0.03$ |
| Cf-SGBP-CBMxx<sub>II/III</sub> | $2.76 (\pm 0.22) \times 10^4$ | $-19.2 \pm 1.11$ | $-45.17 \pm 3.95$ | $0.73 \pm 0.03$ | $-5.95 \pm 0.05$ |
| Cf-SGBP-CBMxx<sub>III/IV</sub> | $3.63 (\pm 0.44) \times 10^4$ | $-18.6 \pm 1.12$ | $-42.45 \pm 4.04$ | $0.76 \pm 0.04$ | $-6.11 \pm 0.06$ |
| Cf-SGBP-CBMxx<sub>IV</sub> | $1.04 (\pm 0.03) \times 10^5$ | $-14.0 \pm 0.12$ | $-24.78 \pm 0.46$ | $0.49 \pm 0.01$ | $-6.73 \pm 0.03$ |
| **Cf-SGBP-CBMxx<sub>IV</sub> - laminarin-derived oligosaccharides (DP7 and DP5)** |  |  |  |  |  |
| | $K_a$<br>[$\text{M}^{-1}$] | $\Delta H$<br>[kcal mol<sup>-1</sup>] | $\Delta S$<br>[cal mol<sup>-1</sup> K<sup>-1</sup>] | N<br>[sites] | $\Delta G$<br>[kcal mol<sup>-1</sup>] |
| Cf-SGBP-CBMxx<sub>IV</sub><br>DP7 | $1.87 (\pm 0.17) \times 10^4$ | $-7.99 \pm 0.02$ | $-7.72 \pm 0.19$ | $0.66 \pm 0.02$ | $-5.73 \pm 0.05$ |
| Cf-SGBP-CBMxx<sub>IV</sub><br>DP5 | $1.39 (\pm 0.07) \times 10^4$ | $-9.14 \pm 0.18$ | $-12.20 \pm 0.70$ | $0.61 \pm 0.04$ | $-5.56 \pm 0.04$ |

### Two elongated multidomain proteins allow for laminarin recognition and scavenging

The multidomain structure of the *C. forsetii* proteins and corresponding ITC experiments for the Cf-SGBP constructs raised the question about the spatial organization of their polypeptide chains. AF predictions were highly uncertain for the linkers connecting the individual modules. Therefore, we analyzed the Cf-SGBP and the Cf-GH16_3 by small-angle X-ray scattering (SAXS), which revealed that both multidomain proteins feature an elongated structural organization (Fig. 4a, Fig. S8, Table S2).

**Fig. 4.**
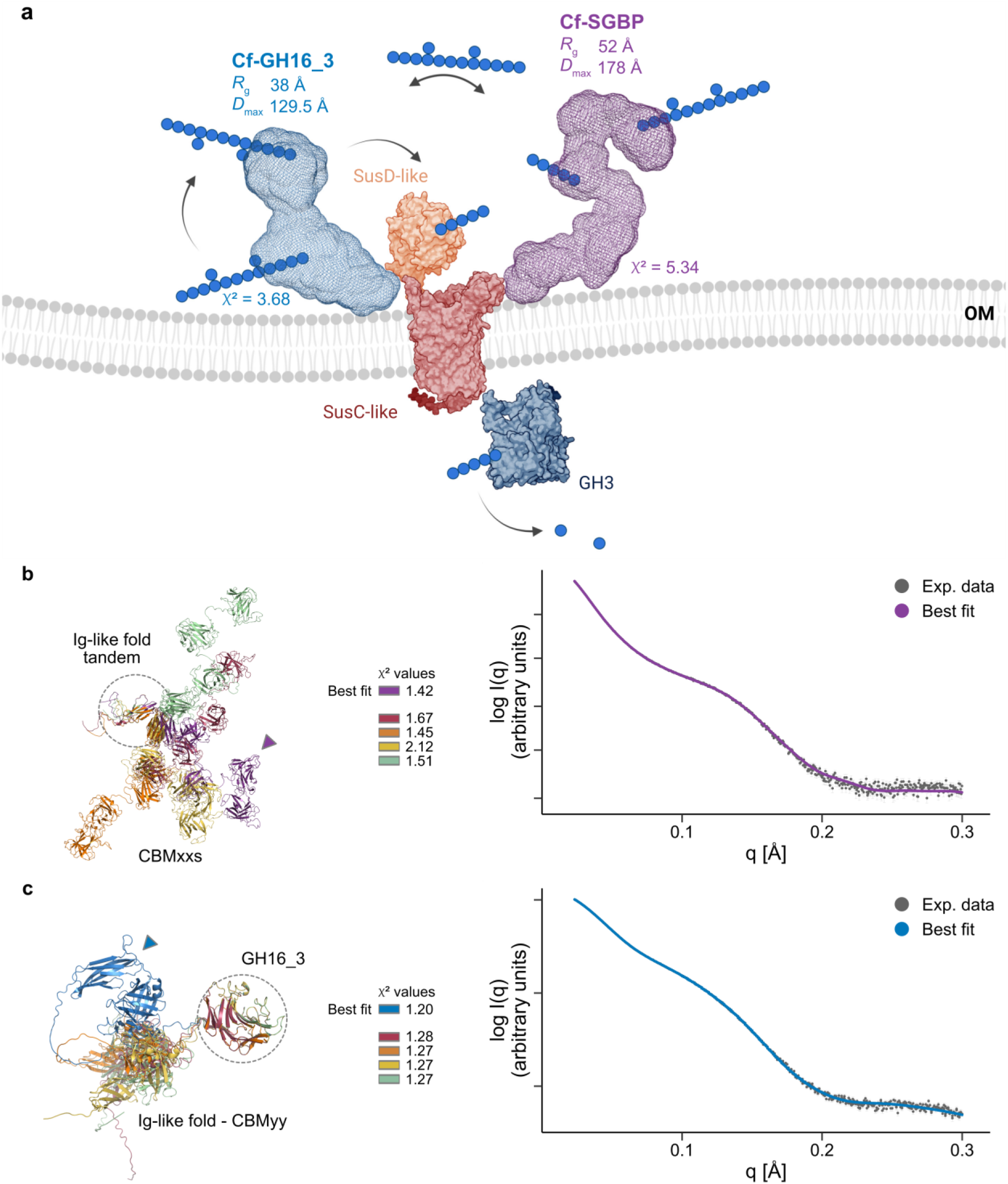
Two elongated multidomain proteins to sense and capture laminarin. **a** *R*_g_ and *D*_max_ values calculated from the experimental SAXS data revealed an elongated structure for the Cf-SGBP and the Cf-GH16_3, which is also reflected in two calculated models for protein envelopes using GASBORi. The figure was created with BioRender using an adapted tentative model of laminarin utilization from (27). The spatial organization of domains in the **b** Cf-SGBP and the **c** Cf-GH16_3 was investigated using DADIMODO. Shown are five models (left graphs) in which domains used for superimposition are circled. The best fit (highest χ^2^) is highlighted with an arrow head and was plotted against the experimental data (right panels). Further information on domains is provided in Fig. 1b.

The values for the radius of gyration (*R*_g_) and the maximum particle distance (*D*_max_) derived from the scattering curve are *R*_g_ 52 Å and *D*_max_ 178 Å for the Cf-SGBP and *R*_g_ 38 Å and *D*_max_ 129.5 Å for the Cf-GH16_3 (Table S2, Fig. S8). The elongated structures were confirmed by the GASBORi (36) *ab initio* calculated envelopes (Fig. 4a, Fig. S9). All calculations for the Cf-GH16_3 returned a bent shape, while much more variable shapes were calculated for the Cf-SGBP (Fig. S9). However, the relatively high χ^2^ values (all >3) of calculated individual shapes for both proteins potentially indicated that the experimental curves were not well fitted by a single envelope, which implicated possible conformational variability. Nevertheless, and using DADIMODO, spatial arrangements of the successive domains based on those predicted by AF could be determined for both the Cf-SGBP and Cf-GH16_3 that individually fitted the experimental curve (Fig. 4b and c, right panels). Superimposition of the individual models, however, covered variable arrangements (Fig. 4b and c, left panels). To further analyze the potential conformational flexibility of these proteins in solution, we used the ensemble optimization method (EOM) (38, 39). Here, an ensemble is selected from a large pool of theoretical conformers, for each of which a corresponding theoretical scattering curve is generated. The selected ensembles for both proteins were able to fit the experimental data, but in contrast to the Cf-GH16_3, the χ^2^ value of the EOM fit of the Cf-SGBP (10.85 *versus* 4.57 for the Cf-GH16_3, Fig. S10) was much higher than χ^2^ values of the GASBORi (Fig. 4a, Fig. S9) or DADIMODO single fits (Fig. 4b and c), indicative of lower conformational flexibility. In addition, the calculated R_flex_ for the selected ensemble of the Cf-SGBP (77%) was lower compared to that of the Cf-GH16_3 (81%), although the R_flex_ of the pool and the final R_Sigma_ were almost the same (Table 2). R_flex_ and R_Sigma_ are values to describe the flexibility in the EOM approach, where 100% or 1 correspond to a fully flexible system, respectively. Finally, a higher flexibility of the Cf-GH16_3 was supported by shifts to higher *R*_g_ and *D*_max_ values for the selected ensemble compared to the pool, whereas they were biased to lower values for the Cf-SGBP (Fig. S11).

**Table 2.** Summary of the EOM-analyses to determine the flexibility of the Cf-SGBP and the Cf-GH16_3 in solution. Given are the final *R*_g_ and *D*_max_ derived from the unique models for the selected ensembles, averaged *R*_g_ and *D*_max_ of the selected ensembles and the pool as well as R_flex_ and R_Sigma_ to describe the flexibility. R_flex_ of 100% or R_sigma_ of 1 corresponds to a fully flexible system.

| | $R_g$ [Å]<br>final | $D_{max}$ [Å]<br>final | $R_g$ [Å]<br>ensemble | $D_{max}$ [Å]<br>ensemble | $R_g$ [Å]<br>pool | $D_{max}$ [Å]<br>pool | $R_{flex}$<br>ensemble/pool | $R_{sigma}$ |
| --- | --- | --- | --- | --- | --- | --- | --- | --- |
| Cf-SGBP | 52.6 | 175.2 | 53.8 | 175.7 | 58.0 | 189.3 | 77%/ 86% | 0.93 |
| Cf-GH16_3 | 37.7 | 127.6 | 37.6 | 126.6 | 35.6 | 118.7 | 81%/ 85% | 0.94 |

### CBMxx and CBMyy are widely distributed in Bacteroidota that thrive during phytoplankton blooms

Screening of bacterial metagenome-assembled genomes (MAGs) obtained from North Sea phytoplankton bloom-associated microbial biomass samples at Helgoland Roads in 2016, 2018 and 2020, yielded 427 sequences coding for CBMxx- and CBMyy-containing protein sequences (555 representative MAGs of which 201 belonged to *Bacteroidota*). The proportion of CBMxx-containing proteins was higher (362) compared to CBMyy (82), whereas only some sequences contained CBMxx and CBMyy (17) (Fig. 5). We found a total of 22 domain combinations, in which CBMxx and/or CBMyy were detected in protein-coding sequences with or without known catalytic modules, e.g., linked to CBM4, CBM6, GH16_3 or rare examples of GH2, GH5_34, GH5_46, GH81 and GH162. The most frequent organizations included CBMxx- or CBMyy-only-proteins (72% for CBMxx with up to eight repeats; 52% for CBMyy, always alone) followed by CBMxx or CBMyy associated with a GH16_3 (20% for CBMxx, 26% for CBMyy). In addition, corresponding sequences were frequently co-localized with genes coding for known laminarin-active enzymes, e.g., GH16_3, GH17 or GH30 family members, but also with *susCD*-like genes. Similar pictures of domain organization and genomic context for both CBMs emerged from screening draft genomes of 53 published North Sea *Bacteroidota* (6), including *C. forsetii*, as well as of the CAZy PULDB for both CBMs (as of May 2023). A total of 92 CBMxx- and/or CBMyy-containing sequences were detected in 38 of the strains from the North Sea (Fig. S12) and >400 over ∼2,000 *Bacteroidota* genomes with additional relevant partners such as GH3, GH149 and GH5_46 using CAZy, strengthening this linkage. In the bloom data set, CBMxx was predominant in *Bacteroidota*, but also occurred in *Proteobacteria* or as rare exceptions in *Actinobacteriota* and *Myxococcota*, whereas CBMyy was exclusive to *Bacteroidota*. This was confirmed by CAZy, which additionally showed that CBMxx and CBMyy occur also in environments other than marine. Structural deviations that we have already discovered in CBMxxs of the Cf-SGBP are also reflected in sequences from bacterial MAGs (Fig. S13 and S14). Nevertheless, residues mediating polar interactions with the substrate in the Cf-SGBP-CBMxx_IV_, and also some structural residues, were highly conserved (Fig. S14). Since not all putative non-catalytic proteins may be attached to the outer membrane to function as CBM-containing SGBPs, we will henceforth use the term (S)GBPs.

**Fig. 5.**
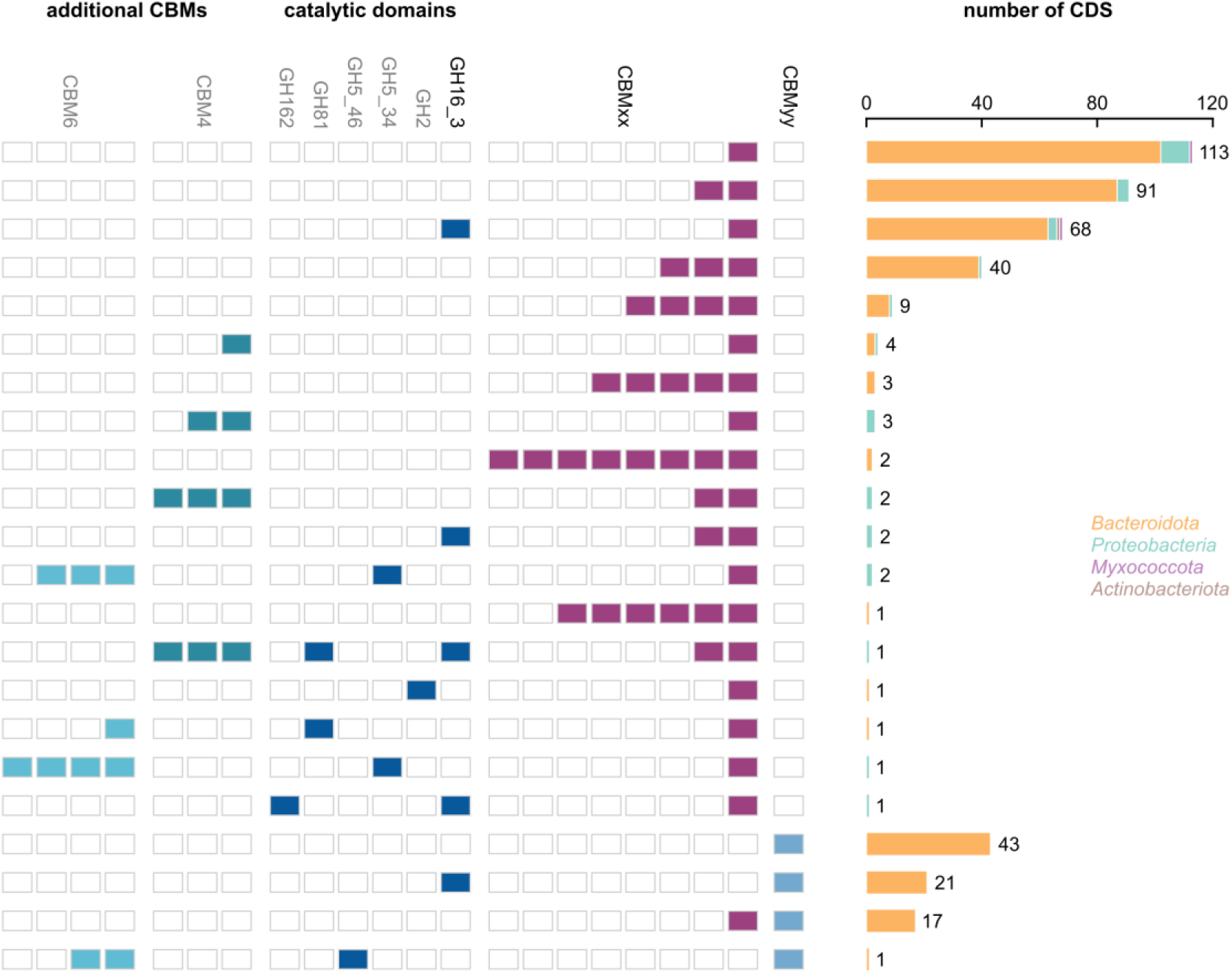
Distribution of CBMxx and CBMyy in proteins encoded in MAGs from algae blooms in the North Sea. Overview of detected CBMxx- and CBMyy-coding sequences and their co-occurrence with CBMs and catalytic modules in multimodular proteins as well as the frequency of occurrence of detected domain combinations and the assigned taxonomy. The graph also depicts multiplicity of certain modules, where each tile represents a detected domain. Note that this graph does not represent the organization of domains within the polypeptide chain. While CBMxx crossed phyla, CBMyy was restricted to *Bacteroidota* in this data set.

### Protein expression correlated with the course of the bloom

Metaproteome analyses showed that the overall expression of CBMxx and CBMyy-containing proteins correlated with bloom intensities (Fig. 6a and b), as assessed by chlorophyll *a* measurements (40). CBM-containing (S)GBPs were more prevalent than CBM-containing enzymes (mainly GH16_3s), possibly due to the higher number of putative (S)GBP sequences. In addition, production of CBM-containing (S)GBPs remained more or less stable after chlorophyll *a* peaks. Looking at the expression of specific laminarin PULs derived from MAGs, CBMxx-containing (S)GBPs, but also CBMyy-containing GH16_3s were detected with a high coverage, while other PUL-encoded GHs were often not captured (e.g., GH3, GH17 or GH30_1). Moreover, CBMxx- and CBMyy-containing proteins were highly abundant in metaproteomes, sometimes comparable to the expression of SusC/D-like ‘marker’ proteins (Fig. 6c).

**Fig. 6.**
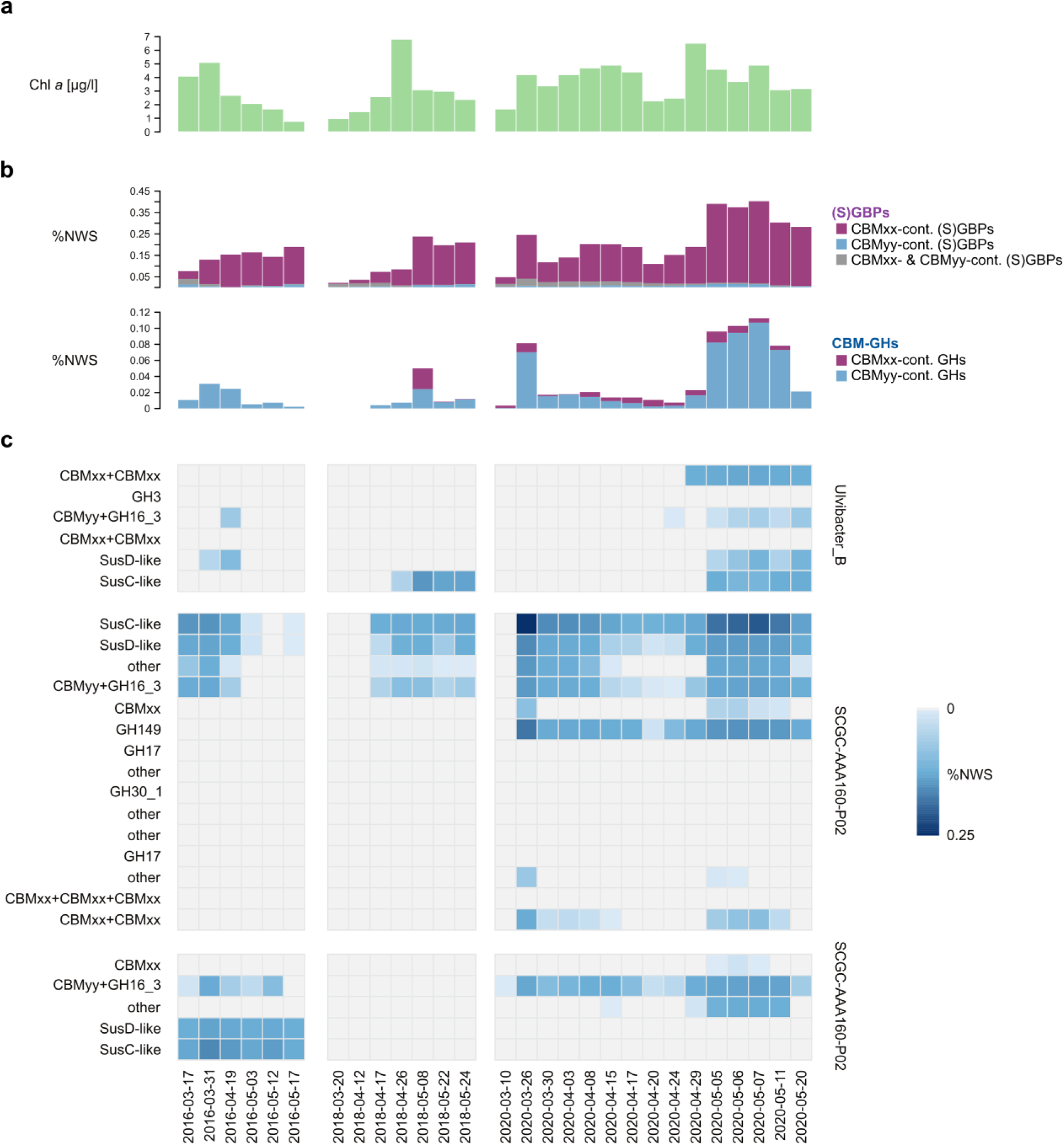
Expression of CBMxx- and CBMyy-containing proteins correlates with the course of phytoplankton blooms. Data show chlorophyll *a* measurements (40) and results of semi-quantitative metaproteome analyses (0.2 µm bacterial fractions) from three respective years, see Material and Methods. **a** Course of the bloom illustrated using the chlorophyll *a* concentration. **b** Summed protein abundance values (%NWS, normalized weighted spectra) for CBMxx- or CBMyy-containing proteins. The upper graph depicts putative (S)GBPs, containing CBMs only, and the lower graph depicts CBM-containing catalytic proteins. Please note different scales (%NWS) for both graphs. **c** Abundance of proteins encoded by three selected laminarin PULs. Gray tiles correspond to proteins that were not quantified.

## Discussion

Elevated photosynthetic primary production during marine phytoplankton blooms lead to pronounced increases in available substrates and nutrients to marine bacteria. These include a plethora of polysaccharides where laminarin is particularly abundant (1). The surface of many bloom-associated bacteria is therefore equipped with specific sugar-binding and -degrading proteins to ensure the recognition and docking of target substrates to the cellular surface, to allow first steps of enzymatic catalysis, but also to prevent loss of small-sized oligosaccharides that need to be shuttled into the cell. The successful and specific binding of poly- and oligosaccharides is therefore a key process in bacterial polysaccharide degradation that can only be understood by detailed analyses of the CBMs involved, as exemplified in this study by the two novel laminarin-binding families CBMxx and CBMyy.

### CBMxx and CBMyy bind laminarin, but belong to different 3D folds

CBMxx and CBMyy both bind to laminarin, but differ substantially in sequence and structure. All four CBMxxs of the Cf-SGBP exhibit a ß-sandwich fold that provides a distinct binding cleft. At the same time, differences in the surrounding loops result in variation of the wideness and depth of the cleft. While CBMxx_I/II/IV_ display a deep and narrow cleft, it is much shallower in CBMxx_III_. This is consistent with observations in other multidomain proteins with multiple CBMs of the same family, e.g., the CBM22 tandem in a xylanase from *Paenibacillus barcinonensis* (41). Here, a wider groove in one of the CBM22 units was suspected to allow for binding of larger decorations. Screening of MAGs of bloom-associated bacteria for CBMxx sequences revealed similar structural variations that may affect either binding specificity or affinity. In contrast to the common ß-sandwich fold of the CBMxx, the ß-barrel fold of the CBMyy is rather rare in CBMs. The latter has recently been described in a CBM-containing SGBP from *B. fluxus,* which can now be classified as a CBMyy. Interestingly, in addition to laminarin, the Cf-GH16_3-CBMyy was also able to bind MLG lichenan, as previously observed for the *B. fluxus* protein (21). Moreover, the Cf-GH16_3-CBMyy lacks a distinct cleft. In the case of the *B. fluxus* protein, binding of ß-glucan termini occurred on the top of the ß-barrel, in a very shallow cleft or platform (21), which we likewise suggest for the *C. forsetii* protein (Fig. S2). This binding site, distally positioned within the loop regions connecting the successive β-strands of the barrel core, is to some extent comparable to that of some CBM6 (42) or CBM32 (43) members, and of several lectin classes (https://unilectin.unige.ch/unilectin3D/). In summary, CBMxx classifies as a type B CBM, probably with more or less possibility to accommodate branches, while CBMyy classifies as a type C CBM (44).

### Shape and flexibility of CBMxx- and CBMyy-containing proteins facilitate recognition and processing of laminarin

The laminarin-binding CBMxx and CBMyy are arranged in elongated shapes of multimodular proteins. In the case of the Cf-SGBP, with its four CBMxxs, the protein may tower out other outer membrane proteins, literally reaching out for its target substrate (Figure 4a). Elongated shapes of multimodular CBM-containing SGBPs have been determined before, e.g., for an SGBP in a heparin/heparan sulfate PUL from *B. thetaiotaomicron* (45). Like the Cf-SGBP (SAXS: *R*_g_ 52 Å and *D*_max_ 178 Å), the *B. thetaiotaomicron* protein consisted of six distinct domains and SAXS determined similar spatial expansions (*R*_g_ of 44.3 Å and *D*_max_ of 150 Å). In addition to the elongated shapes, analyses revealed a bent shape for CBMyy-containing Cf-GH16_3 (*R*_g_ 38 Å and *D*_max_ 129.5 Å), seen before in another multidomain laminarinase from *Thermotoga petrophila* (CBM-GH16-CBM, *R_g_* 40 Å and *D_max_* 130 Å) (46). Our results also indicated that both proteins are flexible to some extent, albeit with a higher flexibility for the Cf-GH16_3. The increased flexibility may be due to missing prolines in the Cf-GH16_3 linkers, as central prolines are suggested to hamper flexibility (45, 47). Such prolines are present in linkers between CBMxxs in the Cf-SGBP. It was speculated that this rigidity enables distance to the membrane and thus an improved exposure of the sugar-binding site (47). This might be further supported by Ig-like fold domains that could act as spacers to the outer membrane (21). Such a projection of SGBPs into the extracellular space has recently been shown for a fructan-specific SGBP from *B. thetaiotaomicron* in a complex with a GH32 and SusC/D-like proteins, called the ‘utilisome’ (48). Compared to the fructan-specific SGBP, which contains one CBM, the laminarin-specific Cf-SGBP from *C. forsetii* provides four CBM docking sites. However, our analyses indicated binding of two laminarin chains at maximum, possibly due to steric inhibition of binding due to the other CBMs. Nevertheless, equipping the cellular surface with SGBPs that contain multiple CBMs increases binding opportunities in 3D space and might optimize the presence of proteins on the bacterial surface as a competitive advantage compared to single CBM-containing SGBPs. Moreover, multiple binding sites in SusE and SusF are known to compensate for decreased binding due to polysaccharide capsules in gut bacteria (49). This may be transferable to the marine counterpart, which is also known to produce exopolysaccharides (50). In the fructan utilisome the sugar chain was held by an accessory tethering side of the GH32 and by the SGBP. The GH32 catalytic activity then cuts between both held ends, which was suggested to facilitate degradation (48). In general, the advantageous effect of CBM-enzyme co-occurrence has largely been demonstrated (e.g., 51, 52). CBMs may also prevent the loss of released degradation products by serving as a short-term cache for cleaved sugars, which are then further digested or translocated to the SusC/D-like transporters (Fig. 4a). At the same time, the affinity of laminarin-derived oligosaccharides to the Cf-SGBP-CBMxx_IV_ was lower, as also observed for a CBM6 from *Z. galactanivorans* Dsij^T^ (23), which supports its major role in capturing laminarin.

### Vivid exchange and combination of domains to increase fitness

Our data from phytoplankton bloom-associated bacteria show that the *C. forsetii* proteins are representatives of diverse domain organizations of both CBMs in multimodular catalytic proteins, mostly GH16_3s, and putative non-catalytic proteins of which a proportion might function as SGBPs. This suggests a rather dynamic intermixing of CBMs by translocations, duplications or losses, which causes also turnovers between catalytic and non-catalytic proteins. Structural variability of CBMs further contributes to this diversity, which may determine substrate specificity and/or affinity. This spectrum of domain combinations and protein structures in sugar-catching and processing proteins likely increases competitiveness to also encounter structurally similar substrates, such as different laminarins or other ß-glucans. The environmental relevance of these novel laminarin-binding CBM families CBMxx and CBMyy was further reflected by a good coverage of corresponding proteins during phytoplankton blooms, which was so far mostly observed for SusC/D-like proteins and GH16_3s associated to laminarin degradation (4, 8). Protein expression of laminarin-binding CBMxx- and CBMyy-containing proteins correlated with the overall intensity of the investigated blooms, supporting their role in marine laminarin use.

In summary, marine bacteria employ CBM-containing enzymes and SGBPs on their cellular surface to scavenge laminarin from the water column. This is facilitated by elongated shapes of these multimodular proteins, where a higher flexibility of the polypeptide chain may increase catalytic activity. In comparison, multiple CBMs and the limited flexibility of SGBPs may allow the accumulation of laminarin, making SGBPs optimal sugar-trapping antenna to further enhance catalysis. Together with the wide distribution and high expression of CBMxx- and CBMyy-containing proteins during phytoplankton blooms, our findings corroborate the key role of the two novel CBMs in laminarin utilization.

## Materials and Methods

### Carbohydrates

Laminarin (*Laminaria digitata*) was acquired from Sigma-Aldrich (Merck kGaA, Darmstadt, Germany) and MLG lichenan (Icelandic Moss) from Megazyme International Ltd. (Bray, Ireland). Laminarin-derived oligosaccharides were produced following the protocol from Jam *et al.* (23) using a GH16_3 (LamA) from *Zobellia galactanivorans* Dsij^T^ (13).

### Nomenclature

We refer to the full-length surface glycan-binding protein, which is SusE-positioned in the laminarin PUL of *Christiangramia forsetii* KT0803^T^, as Cf-SGBP (locus tag GFO_RS17395). The PUL further encodes two GH16_3s. Subject of this study was the GH16_3 encoded downstream of the Cf-SGBP, which we refer to as Cf-GH16_3 (locus tag GFO_RS16360). The two proteins display a multidomain structure and specific domains were further specified, e.g., Cf-GH16_3-Cat refers to the catalytic module of the Cf-GH16_3. In the case of repetitive domains of the Cf-SGBP, these modules were enumerated either alphabetically for Ig-like fold domains or with Roman numerals for the CBMxxs, starting from the N-terminus and indicated as subscripts, e.g., Cf-SGBP-Ig_a_ refers to the first of two N-terminal Ig-like fold domains of the Cf-SGBP. Whenever we refer to an SGBP in the text, we refer to CBM-containing SGBPs, excluding SusD-like proteins.

### Cloning and site-directed mutagenesis

We used Alphafold2 (28) and Phyre2 (29) to spatially define individual modules of the two multidomain proteins. LipoP (53) detected signal peptides and MeDor (54) predicted protein disorder by hydrophobic cluster analysis. In combination, these tools served to define suitable cloning sites to express full-length proteins, individual domains or combinations thereof. Corresponding genes were amplified from *C. forsetii* genomic DNA by PCR using primers listed in Table S3. The PCR products were cloned into expression vectors pRF3 (ampicillin resistance) or pET28a (kanamycin resistance) by restriction-ligation to produce recombinant N-terminal hexa-histidine tagged proteins. The vector pRF3 is a derivative of pFO4 (55). The full-length *Cf-SGBP* and *Cf-GH16_3* were cloned into pRF3. PCR-products were digested using BamHI and AvrII, and the pRF3 vector using BamHI and NheI. In all other cases, PCR products and the pET28a vector were digested using NdeI and SacI. Digested PCR-products were ligated into expression vectors using T4 DNA ligase. Corresponding plasmids for Cf-SGBP-Ig_a/b_, Cf-SGBP-CBMxx_I_, Cf-GH16_3-Ig and Cf-GH16_3-CBMyy in pET28a, including NdeI and SacI restriction sites, were designed and ordered from GenScript (GenScript Biotech, Netherlands).

In addition, we generated two mutants of the Cf-GH16_3-Cat, where each catalytic glutamic acid was replaced by a serine, Cf-GH16_3-Cat_E442S_ and Cf-GH16_3-Cat_E447S_. This was achieved in two steps: first, two separate reactions of PCR, using the Cf-GH16_Cat-encoding plasmid as a template, were carried out to produce the mutant gene as two overlapping fragments (F1 and F2). The purified PCR products F1 and F2 were then used as a template for a final PCR to produce the fused product. To produce the overlapping fragments, we used a forward primer for the first fragment (F1), which annealed to the vector backbone directly upstream of *Cf-GH16_3-Cat*, and a reverse primer for the second fragment (F2), which annealed to the vector backbone downstream of *Cf-GH16_3-Cat* (designated as vector primers, Table S3). These primers were combined with a forward (F2) or reverse (F1) primer that contained the corresponding desired mutation and that annealed to the intended region of the mutation (Table S3). The purified PCR products F1 and F2 were digested with DpnI to remove any residual parental DNA before a final PCR fused F1 and F2 using the vector primers. PCR products were digested with NdeI and SacI and cloned into pET28a as described before.

Sequence identity for all constructs and successful mutation was verified by sequencing (Eurofins Genomics, Ebersberg, Germany). Plasmids were transformed into propagation strains *Escherichia coli* DH10B or DH5α before being transformed into the expression strain *E. coli* BL21(DE3).

### Protein overexpression and purification

The Cf-SGBP and the Cf-GH16_3 were produced in ZYP-5052 auto-induction medium (56), supplemented with 100 µg ml^-1^ ampicillin, for 72 h at 20 °C and 130 rpm. For the production of the other proteins, clones were cultured in LB, supplemented with 30 µg ml^-1^ kanamycin, at 37°C and 130 rpm until IPTG induction (1 mM) and then at 20 °C and 130 rpm overnight. Cells were harvested by centrifugation and pellets were stored at -20 °C until protein purification. Cells were disrupted by chemical lysis. In brief, the cell pellet was suspended in resuspension buffer (50 mM MOPS pH 8.0, 25% sucrose, lysozyme) and agitated for 15 min at 4°C. Then, double the volume of lysis buffer (20 mM MOPS pH 7.5, 100 mM NaCl, 1% sodium DCA, 1% Triton-X) was added, followed by another incubation of 5 min at 4°C. MgCl_2_ concentration was adjusted to 5 mM and samples were incubated with DNase at room temperature until viscosity of the sample decreased. Cell debris was removed from protein extract by centrifugation. Protein extract was loaded onto Histrap HP columns (Cytiva, Vélizy-Villacoublay, France) equilibrated in buffer A (200 mM NaCl, 10 mM MOPS pH 7.8, 20 mM imidazole) and charged with 0.1 M NiSO_4_. In general, 1 ml columns were used for a culture volume of 250 ml and 5 ml columns for 1 l of culture. Proteins were eluted with a linear gradient of 0-100% buffer B (200 mM NaCl, 10 mM MOPS pH 7.8, 500 mM imidazole) within 30 min (1 ml column) or 60 min (5 ml column) at a flow rate of 1 ml min^-1^. For SAXS analyses, crystallography or in case of impurities, the quality of the protein sample was further increased by size exclusion chromatography (SEC) using a Superdex 200 16/60 column (Cytiva) in buffer C (200 mM NaCl, 10 mM MOPS pH 7.8). Otherwise, samples were desalted on a Hiprep 26/10 desalting column (Cytiva) in buffer C. Buffers used to purify the proteins for SAXS analyses contained 100 mM NaCl. Samples were evaluated by SDS-PAGE. Protein concentration was determined on a Nanodrop Spectrophotometer using the molecular weight and the extinction coefficient. If necessary, samples were concentrated at 3,500 rpm and 4°C using Amicon Ultra centrifugal units (Merck).

### Affinity gel electrophoresis (AGE)

Prior to AGE, concentration of NaCl was decreased from 200 mM to 20 mM by diluting the protein sample with 10 mM MOPS (pH 7.8). Recombinant proteins and BSA, which served as a non-binding control, were loaded on 12% native gels without the addition of any carbohydrate, or supplemented with 0.2% laminarin or 0.1% lichenan. Runs were performed at 80 V and 4°C on ice. Proteins were stained using Coomassie Blue. Differences in length of running fronts were considered to evaluate binding.

### Enzyme activity

3,5-dinitrosalicylic acid (DNS) reducing sugar assay (57) was used to confirm activity of the Cf-GH16_3 on laminarin and MLG lichenan. Proteins (20 mM NaCl, 10 mM MOPS pH 7.8) were mixed with 0.5% laminarin or lichenan (20 mM NaCl, 10 mM MOPS pH 7.8) in 200 µl reactions. Samples were incubated overnight at room temperature and therefore experiments correspond to endpoint measurements. The non-catalytic Cf-SGBP and samples without the addition of protein served as controls. 100 µl of DNS reagent was added to 100 µl of sample. After 5 min of incubation at 95 °C, the reaction was stopped by adding 800 µl of water. Absorbance was measured at 540 nm and increased with catalytic activity. Experiments were performed in triplicates. Absorbance did not increase in the Cf-GH16_3 incubations with MLG lichenan. In this case, samples were further investigated by fluorophore-assisted carbohydrate electrophoresis (FACE) that allows for higher sensitivity. 50 µl of the overnight incubations were dried under vacuum and mixed with 2 µl of 0.15 M 8-aminonaphtalene-1,3,6-trisulfonate (ANTS) and 2 µl of 1 M NaBH_3_CN. Samples were incubated at 37 °C in the dark for at least 2 h and dried under vacuum. After resuspension in 20 µl of 25% glycerol, 5 µl were loaded onto 27% acrylamide gels and migrated at 200 V and 4 °C. Results were visualized under UV.

### Isothermal titration calorimetry (ITC)

Proteins were dialyzed against 100 mM NaCl, 10 mM MOPS pH 7.8. The dialysis buffer was used to prepare laminarin and laminarin-derived oligosaccharide solutions, as well as for washing of the ITC cell and for controls. Prior to measurements, samples and solutions were centrifuged for degassing and removal of putative protein aggregates. In addition, protein samples were evaluated by SDS-PAGE and Nanodrop repeatedly. ITC was done on a MicroCal^TM^ ITC 200 machine. Laminarin or laminarin-derived oligosaccharides were injected into 200 µl of protein, which was loaded into the sample cell. 2.5 mM of laminarin was injected into 147.6 µM of the Cf-SGBP, into 127.7 µM of the Cf-SGBP-CBMxx_II/III_ or into 94.7 µM of the Cf-SGBP-CBMxx_III/IV_. 1 mM of laminarin was injected into 147.5 µM of the Cf-SGBP-CBMxx_IV_. For calculations with laminarin, we used the average molecular weight of laminarin from *Laminaria digitata* (2). In addition, 5 mM of laminariheptaose (DP7), 10 mM of laminaripentaose (DP5) and up to 20 mM of laminaribiose (DP2) were injected into 359.6 µM of the Cf-SGBP-CBMxx_IV_. Settings were as follows: cell temperature 20 °C (293.15 K), reference power 10 µCal/s, stirring speed 750 rpm, filter period 1 sec, injection spacing 300 sec, 1 µl injection volume (first injection 0.3 µl), 35 injections in total. Experiments were performed in at least triplicates. 2.5 mM and 5 mM of laminarin were injected into 118.8 µM of the Cf-SGBP-Ig_a_ as a non-interacting control. Titrations of buffer into proteins, laminarin into buffer or buffer into buffer represented additional controls. MICROCAL ORIGIN v. 7 was used to analyze the data. A single-site binding model was selected.

### SEC-SAXS

SAXS data of the Cf-SGBP and the Cf-GH16_3 were collected at the Synchrotron SOLEIL on the SWING beamline. Protein samples were centrifuged prior to analyses to remove putative aggregates. SAXS measurements were coupled with prefixed SEC. The SEC column was equilibrated in buffer C (100 mM NaCl, 10 mM MOPS pH 7.8), which was used for protein purification. 70 µl of protein were injected, the Cf-SGBP at 8.2 mg ml^-1^ and the Cf-GH16_3 at 12.9 mg ml^-1^. The sample-detector-distance was 2.4 m, resulting in a scattering vector *q*-range of 0.012 – 0.504 Å^-1^ for the Cf-SGBP and 0.017 – 0.504 Å^-1^ for the Cf-GH16_3. The obtained scattering data were normalized and corrected according to standard procedures. The Guinier equation was used to calculate the forward scattering *I*(*0*) and the radius of gyration *R*_g_. The distance distribution function *P(r)* and the maximum particle dimension *D*_max_ were calculated by Fourier inversion of the scattering intensity *I(q)* using GNOM integrated in the PRIMUS software (ATSAS 3.1.3) (58). Models of protein envelopes were calculated from the experimental scattering curves using GASBORi (36). High χ^2^ values indicated flexible proteins. Therefore, the data were analyzed using DADIMODO (37). Here, structural data of proteins are included into the analysis to identify an arrangement that best describes the experimental data. For this, rigid bodies and linkers were defined based on the structures predicted by Alphafold2 (28). Finally, EOM analysis was used to describe the flexibility of proteins in solution (38, 39), again using Alphafold2-derived coordinates. Experimental and processed data were visualized in R using the ggplot2 package.

### Crystallography

Crystallization for both recombinant constructs of Cf-SGBP-CBMxx_IV_ (at a concentration of 11.7 mg ml^-1^) and Cf-SGBP-CBMxx_III/IV_ (at a concentration of 9.5 mg ml^-1^) were tempted, in presence or absence of laminarin oligosaccharides (DP3/4), but never led to crystals in the case of Cf-SGBP-CBMxx_III/IV_. Crystallization screening was undertaken with the nanodrop-robot Crystal Gryphon (Art Robbins instruments) starting with sparse matrix sampling kits JCSGplus, PACT, PEGs, and Wizard Classic I/II and Ligand-Friendly Screen (from Qiagen and Molecular Dimensions). The initial crystallization conditions were manually optimized and suitable crystals of Cf-SGBP-CBMxx_IV_ in presence of laminarin-trisaccharides were obtained using the hanging drop vapor diffusion method as follows: 2 µl of protein were mixed with 1 μl of reservoir solution (500 µl) containing 2.5 M ammonium sulfate, 0.1 M BIS-TRIS propane pH 7.0, 6% ethanol and 4% polyethylenglycol (PEG) 6000. Diffraction data were collected on PROXIMA1 (SOLEIL synchrotron) with a single crystal of Cf-SGBP-CBMxx_IV_ in complex with laminarin-trisaccharide at 1.9 Å resolution. The data were integrated using XDS (59) and scaled with aimless (60). The structure was solved by molecular replacement with Phaser (61) using the AF model as starting point. The crystal structure was further enhanced with alternating cycles of refinement with REFMAC (62) and manual construction using COOT (33). All further data collection conditions and refinement statistics are given in Table S1.

### Phytoplankton bloom data

Chlorophyll *a* concentration as well as metagenomes, metagenome-assembled genomes and metaproteomes were obtained from samples taken during spring phytoplankton blooms at North Sea island of Helgoland in 2016 (9), 2018 and 2020 (10, 40). Chlorophyll *a* data were obtained from the Helgoland Roads time series (63) and have been published for the respective years (40). Metagenome sequencing, assembly and binning (MAGs) are described in (9) (2016) and (10) (2020). Preparation of metaproteome samples (0.2 µm fraction) as well as corresponding LC-MS/MS measurements and data analyses have been described in detail in (9) (2016) and in (40) (2018 and 2020).

### Comparative genomics

MAGs from 2016, 2018 and 2020 were dereplicated using dRep v1.14.6 (64) to prevent redundancy, applying a minimum completeness of 70% and contamination lower than 5% at 0.95 ANI (average nucleotide identity). For each representative MAG (555), protein coding sequences for representative MAGs were predicted with Prokka v1.14.6 using default settings (65). PULs, CAZymes, CBMs, SusC/D-like proteins were predicted as described previously (9), using hmmscan v3.3.2 against dbCAN-HMMdb-V1 and DIAMOND BLASTp v2.1.1.155 (66) against CAZyDB.08062022 provided by dbCAN (67). CBMxx- and CBMyy-containing sequences within representative MAGs and additional draft genomes of previously published 53 North Sea *Bacteroidota* (6) were identified using hmmscan as described above, adding corresponding HMM profiles to the database. Results were further filtered using the hmmscan-parser.sh script from dbCAN with an e-value cutoff of 1E-15 and a minimum coverage of 35%. The distribution of domain organization was visualized with UpSetR (68, 69). The official CAZy classification was applied to semi-manually determine the CBM boundaries, the multimodular compositions, the taxonomic distributions in CAZy and the PUL prevalence and partnerships in PULDB.

## Data Availability

The metaproteome datasets for 2018 and 2020, the PDB identifier and SAXS data will be released upon acceptance of the manuscript in a peer-reviewed journal. Metagenome assemblies and MAGs from 2016, 2018 and 2020 phytoplankton blooms are available in the European Nucleotide Archive under the project accessions PRJEB28156 (2016), PRJEB38290 (2018) and PRJEB52999 (2020). Metaproteome data from the 2016 blooms are available in the PRIDE archive (70) through the identifier PXD019294. Phytoplankton data are available on request via Pangaea (https://doi.pangaea.de/10.1594/PANGAEA.864676).

## Acknowledgements

This study was supported by the German Research Foundation (DFG) within the research unit FOR 2406 ‘Proteogenomics of Marine Polysaccharide Utilization’ (POMPU) by grants to Hanno Teeling (TE 813/2–3), Dörte Becher (BE 3869/4-3) and Thomas Schweder (SCHW 595/10-3, SCHW 595/11-3). This work also benefited from the support of the Centre National de la Recherche Scientifique (CNRS) and Sorbonne University (Paris). The authors strongly appreciated to have access to the CristalO platform (FR2424, Station Biologique de Roscoff), which is part of the core facility networks Biogenouest and EMBRC-France. We are grateful for the access to the small-angle X-ray scattering beamline and MX-beamline PROXIMA1 at synchrotron SOLEIL (Saint Aubin, France) and thank the staff for their support on site, especially Aurélien Thureau and Pierre Legrand. We thank the team of the Helgoland sampling campaign, especially Lilly Franzmeyer, and the Biological Station Helgoland (Alfred-Wegener-Institut Helmholtz-Zentrum für Polar-und Meeresforschung, BAH-AWI, Germany), especially Inga Kirstein and Karen Wiltshire. The Helgoland Time Series of the Alfred Wegener Institute is supported by the Helmholtz Association as an LK-II performance category program. We thank Jana Matulla, Jasna Nikolic and Lionel Cladière for support in the lab and Stephanie Markert for helpful comments. MKZ gratefully acknowledges the DFG (SCHW 595/10-3), the PROCOPE Mobility Grant and the ERASMUS Staff Mobility for Training, all of which enabled several research stays in Roscoff.

## Author contributions

MC and MKZ directed the study with contribution from EFB and TS. MKZ wrote the first draft of the paper with writing contributions from MC and EFB. All co-authors provided their input on the manuscript and helped with editing. Figures were created by MKZ, MC and DBa. MKZ and MC performed computational sequence analyses of *C. forsetii* proteins. MKZ performed cloning, protein purification as well as experiments on catalytic activity and protein-sugar interactions. NW and RL supported cloning. EFB, MC and MJ supported experiments on protein-sugar interactions. MC and MKZ performed SAXS analyses and data treatment. AJ and MC performed protein crystallography and MC solved the 3D crystal structure. MJ and DJ produced and purified laminarin-derived oligosaccharides. MJ, TE and FT supported protein quality controls. NT classified CBMxx and CBMyy. DBa and NT performed comparative sequence analyses with support of MC and MKZ. DBa analyzed the phytoplankton bloom data with support of FW, HT, ATS and DBe. All co-authors approved the submitted manuscript.

