## Supplementary Information for "Novel laminarin-binding CBMs in multimodular proteins of marine *Bacteroidota* feature prominently in phytoplankton blooms"

^*^corresponding authors

**The Supporting Information file contains:**

Supplementary Tables S1-S3

Supplementary Figure S1-S14

References

**Table S1**: Data collection and refinement statistics for the crystal structure of the CBMxx_IV_ isolated domain of the Cf-SGBP.

|  | **Cf-SGBP-CBMxx_IV_** |
| --- | --- |
| **Data collection** |  |
| Beamline | PROXIMA 1 |
| Wavelength | 0.98 Å |
| Space group | P2_1_2_1_2_1_ |
| Unit cell | a=55.44 Å; b=58.12 Å; c=91.19 Å |
| Resolution range ^a^ (Å) | 47.42-1.90 (1.95-1.90) |
| Total data | 308298 |
| Unique data | 23977 |
| Completeness (%) | 99.8 (97.2) |
| Mean I/σ(I) | 10.9 (1.3) |
| CC(1/2) | 0.99 (0.58) |
| ^b^R_merge_; ^c^R_pim_(%) | 14.8 (181.3); 4.3 (58.7) |
| Redundancy | 12.9 (10.2) |
| **Refinement statistics** |  |
| Resolution range | 47.42-1.90 (1.95-1.90) |
| Unique reflexions | 23914 (1216) |
| Reflexions R_free_ | 1216 (68) |
| R / R_free_ (%) | 23.0/27.9 (39.5/40.,3) |
| RMSD bond lengths | 0.009 Å |
| RMSD bond angles | 1.61° |
| Overall B factor (Å²) | 34.8 |
| B factor: Molecule A (Å²) | 35.2 |
| B factor: Molecule B (Å²) | 34.8 |
| B factor: Solvent (Å²) | 33.6 |
| B factor: Ligands (Å²) | 36.6 |
| ^a^Values in parentheses concern the high-resolution shell.  ^b^R_merge_ = Σ \| I-I_av_ \| / Σ\| I \|, where the summation is over all symmetry-equivalent reflections.  ^c^R_pim_ = corresponds to the multiplicity weighted R_merge_. | |

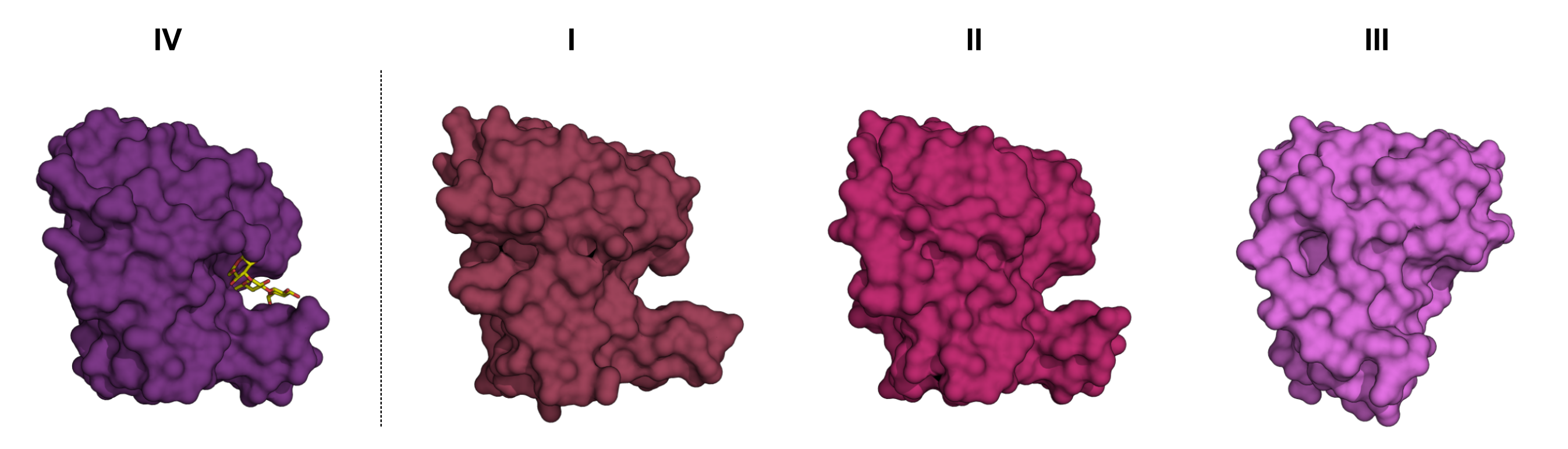

**Fig. S1 Surfaces of the CBMxxs of the Cf-SGBP.** Surfaces of the Cf-SGBP-CBMxx_IV_ and of the Cf-SGBP-CBMxxs_I-III_ derived from the 3D crystal structure or as predicted with Alphafold2, respectively. While modules I, II and IV show quite similar surfaces characterized by a deep and narrow binding cleft, module III displays a very shallow and wider cleft due to a missing loop, see main text.

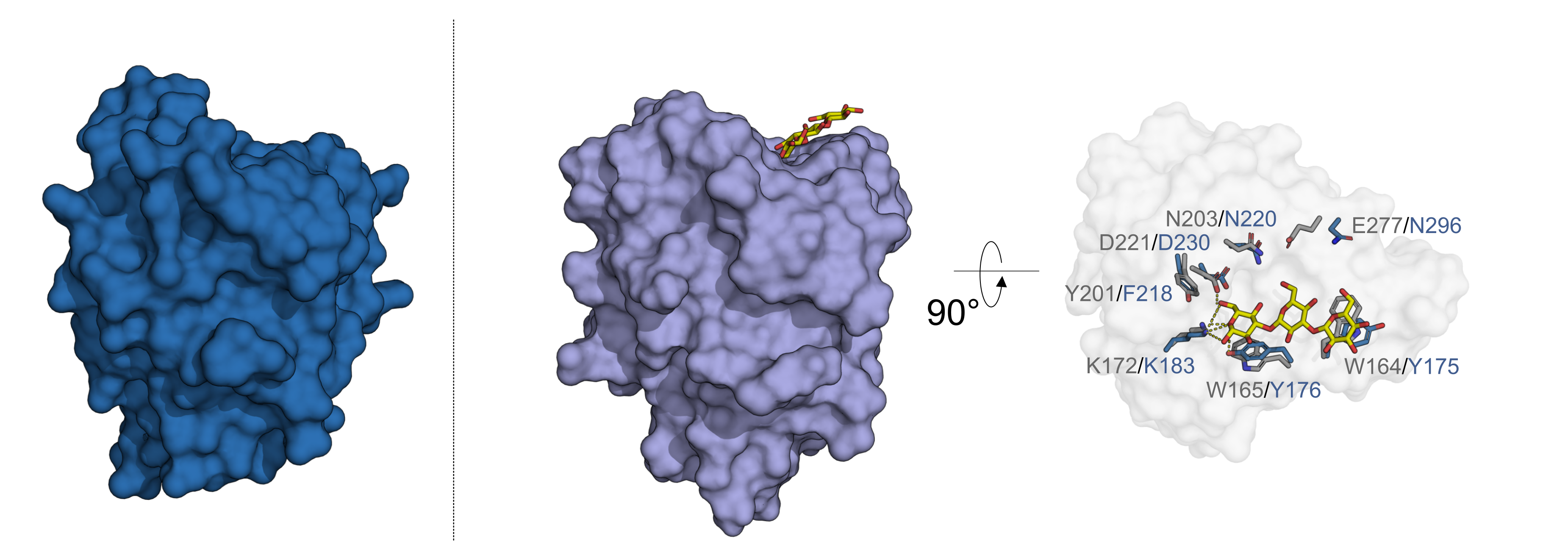

**Fig. S2 The Cf-GH16_3-CBMyy superimposes with a CBM of a SGBP from *Bacteroidetes fluxus*.** Alphafold2 predictions suggest a binding platform for the Cf-GH16_3-CBMyy (blue, left graph) similar to a corresponding laminarin-binding CBM of a SGBP from *B. fluxus* (lightblue, PDB ID 7KV7) (1), see also Fig. 2e in the main text. Top view of the binding platform of the *B.fluxus* CBM shows residues (gray) interacting with laminaritriose and their overlay with residues of the Cf-GH16_3-CBMyy (blue).

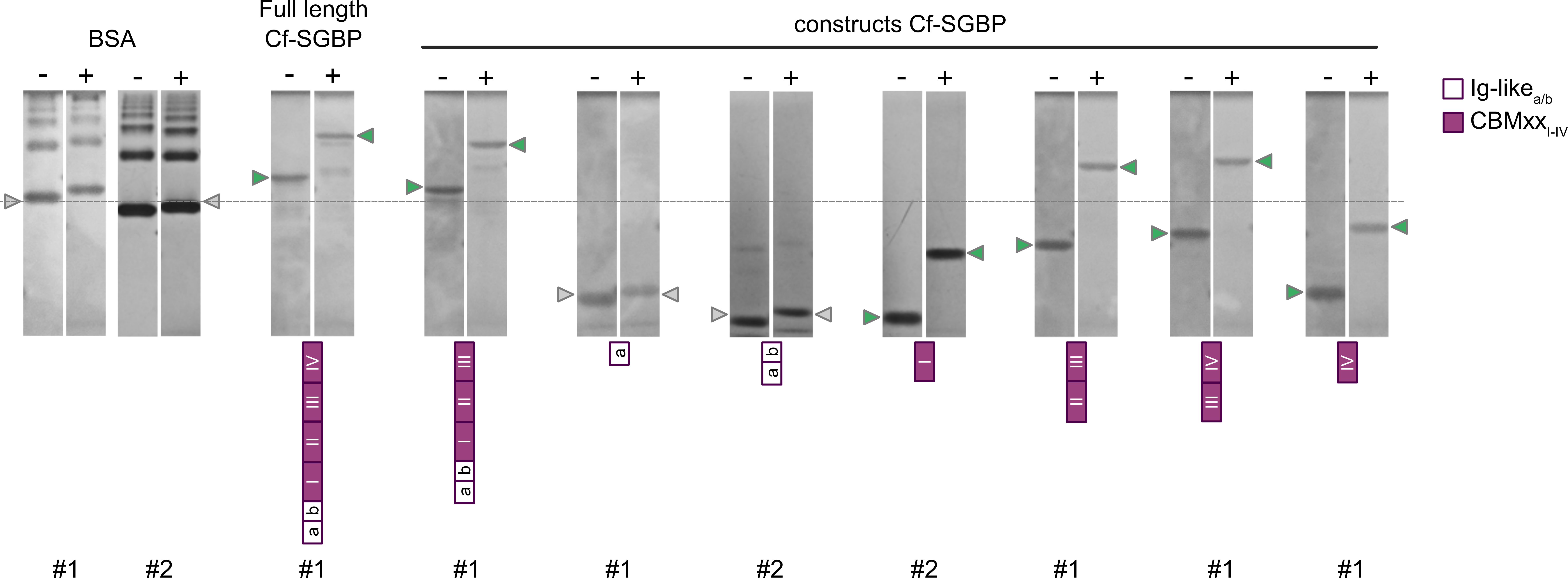

**Fig. S3. The CBMxxs of the Cf-SGBP are binding laminarin.** AGE of the full-length Cf-SGBP as well as of a truncated variant and of dissected domains or domain tandems in the absence (-) or presence (+) of laminarin (0.2%). BSA served as a non-binding control. Green arrow heads indicate proteins considered positive for binding. Two AGE runs were performed separated in time, indicated as runs #1 and #2. Running fronts were considered to evaluate binding.

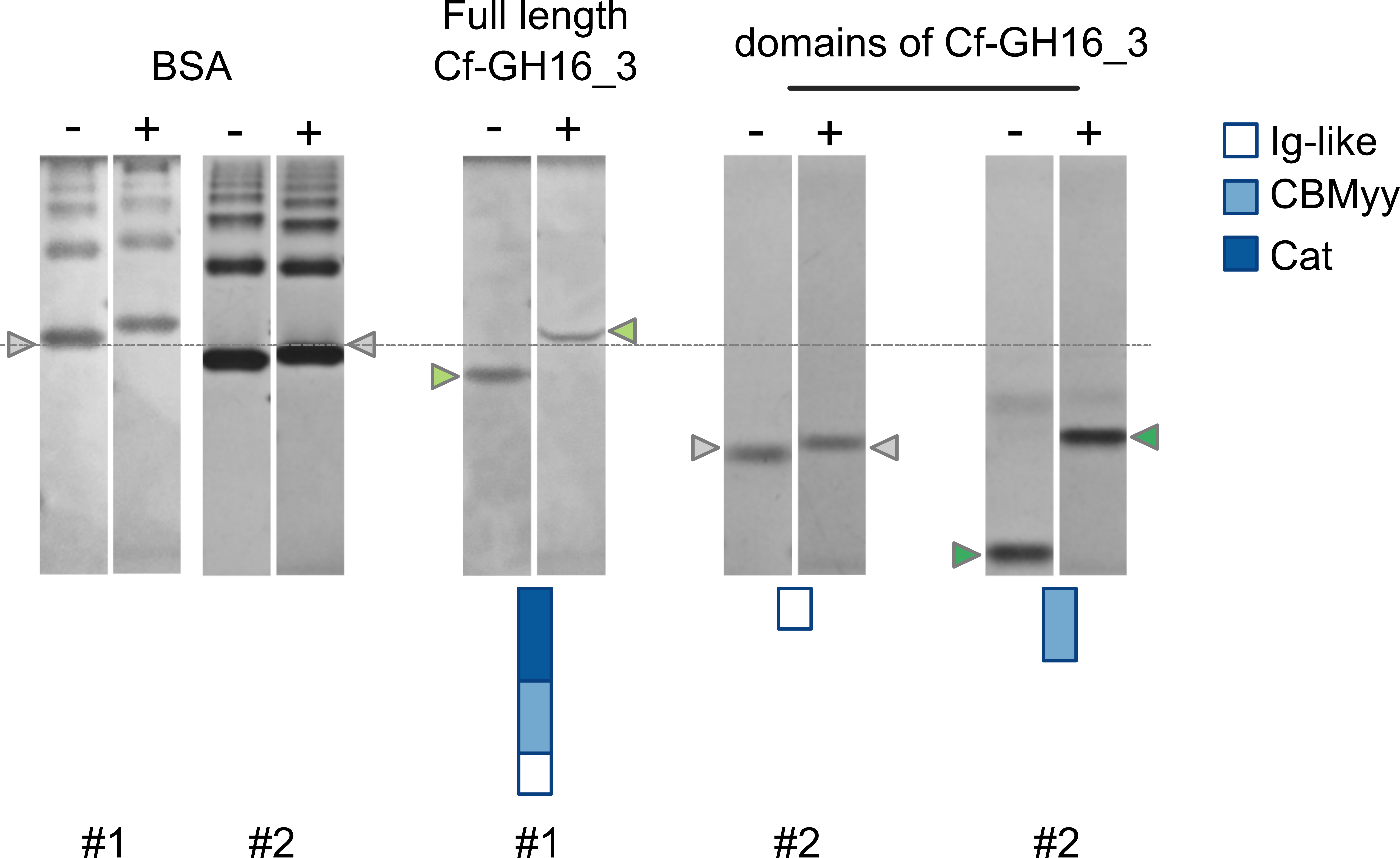

**Fig. S4. The CBMyy of the Cf-GH16_3 binds laminarin.** AGE of the full-length Cf-GH16_3 as well as of two dissected domains thereof, Cf-GH16_3-Ig and Cf-GH16_3-CBMyy, in the absence (-) or presence (+) of laminarin (0.2%). BSA served as a non-binding control. For AGE of the catalytic domain Cf-GH16_3-Cat and mutants thereof, see main text Fig. 3. Green arrow heads indicate proteins considered positive for binding. Retardation of the Cf-GH16_3 is not as clear as for the Cf-GH16_3-CBMyy, which is due to the catalytic activity of the full-length protein, see main text. Two AGE runs were performed separated in time, indicated as runs #1 and #2. Running fronts were considered to evaluate binding.

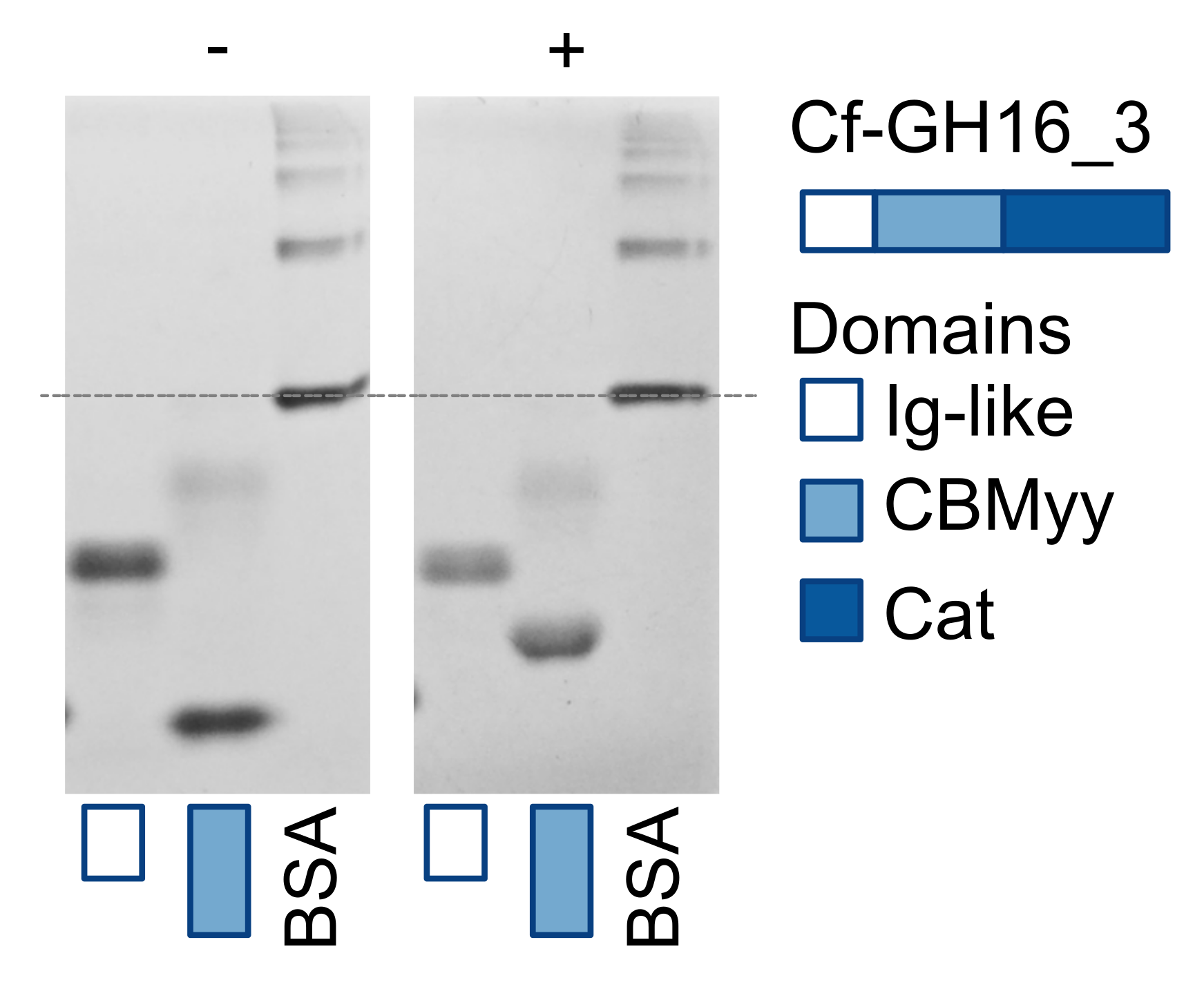

**Fig. S5. The CBMyy of the Cf-GH16_3 is binding lichenan.** AGE of two dissected domains of the Cf-GH16_3 in the absence (-) or presence (+) of 0.1% lichenan. BSA served as a non-binding control. Cf-GH16_3-CBMyy was hold back by lichenan.

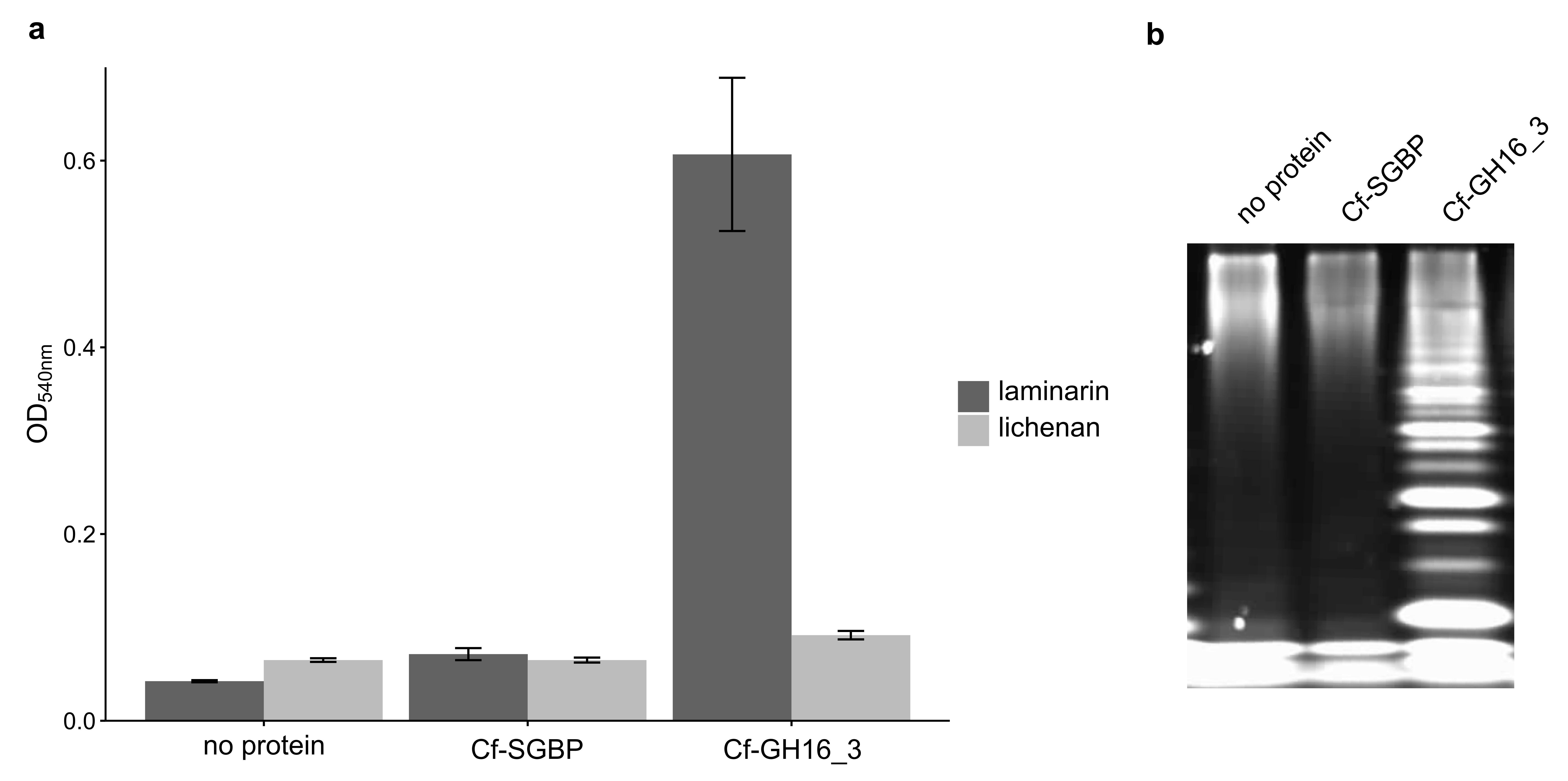

**Fig. S6. The Cf-GH16_3 has a low activity on lichenan.** **a** Activity of the Cf-GH16_3 on laminarin and lichenan as determined by DNS-reducing sugar assay (RSA). The graph represents an endpoint measurement and the OD_540nm_ correlates with the increase of reducing ends during degradation of polysaccharides. Cf-SGBP and samples without protein were used as controls. Experiments were performed in triplicate (n=3). **b** While RSA was not sensitive enough to detect activity of the Cf-GH16_3 on lichenan, activity was confirmed by FACE using the same samples.

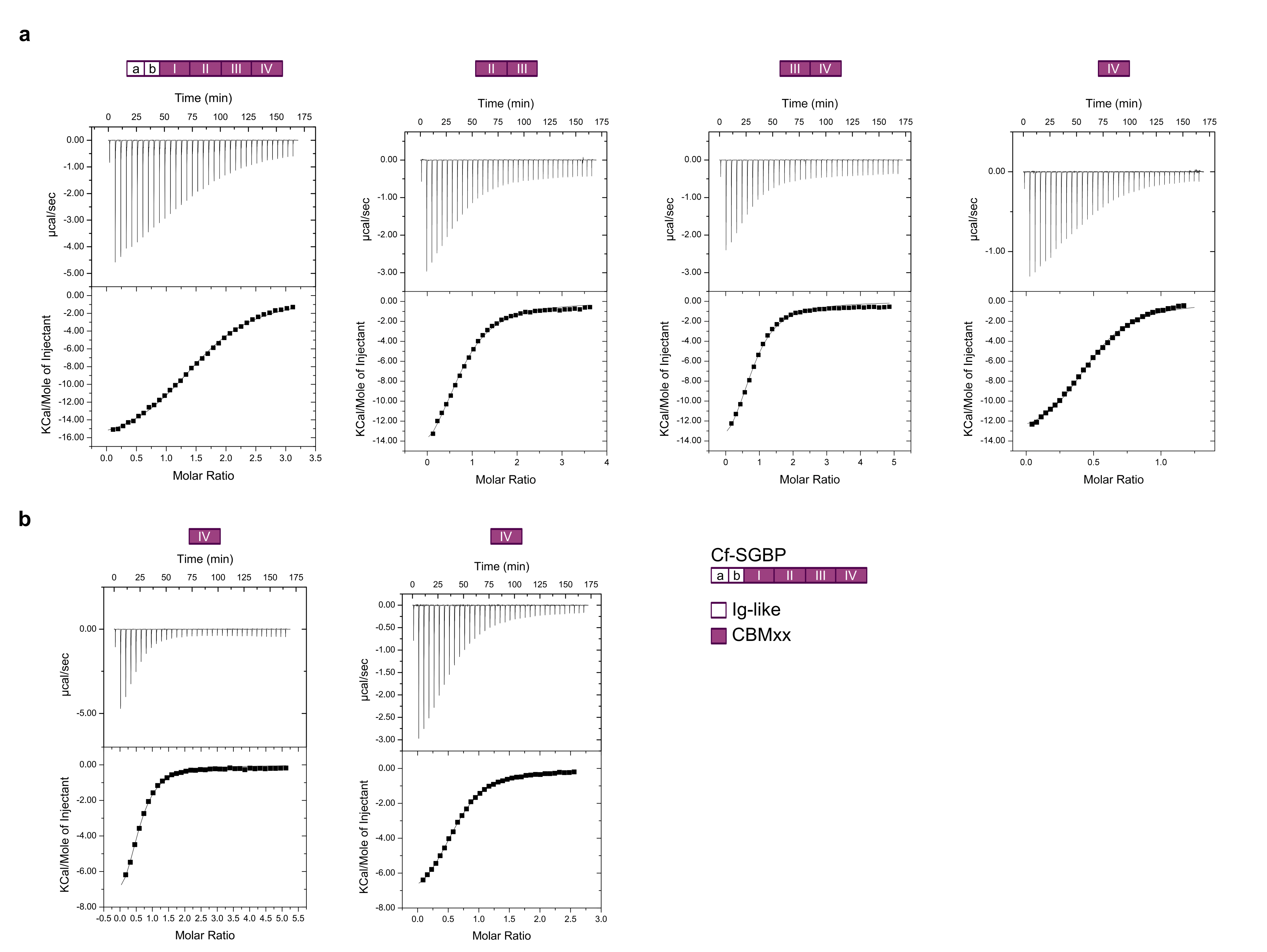

**Fig. S7. Isothermal titration calorimetry (ITC) experiments.** ITC confirmed binding of **a** laminarin to the Cf-SGBP, to the tandems Cf-SGBP-CBMxx_II/III_ and Cf-SGBP-CBMxx_III/IV_ as well as to the single module Cf-SGBP-CBMxx_IV_. **b** ITC also confirmed binding of laminarin-derived oligosaccharides (DP5, left, and DP7, right) to the Cf-SGBP-CBMxx_IV_. The upper panels show heats released upon titrating laminarin or oligosaccharides into the respective protein, whereas the lower panels plot the integrated heat peaks against the molar ratio of ligand. For each protein, one representative experiment of at least triplicates is shown. Corresponding values, derived from the plots are summarized as mean values with standard deviation in the main text, Table 1.

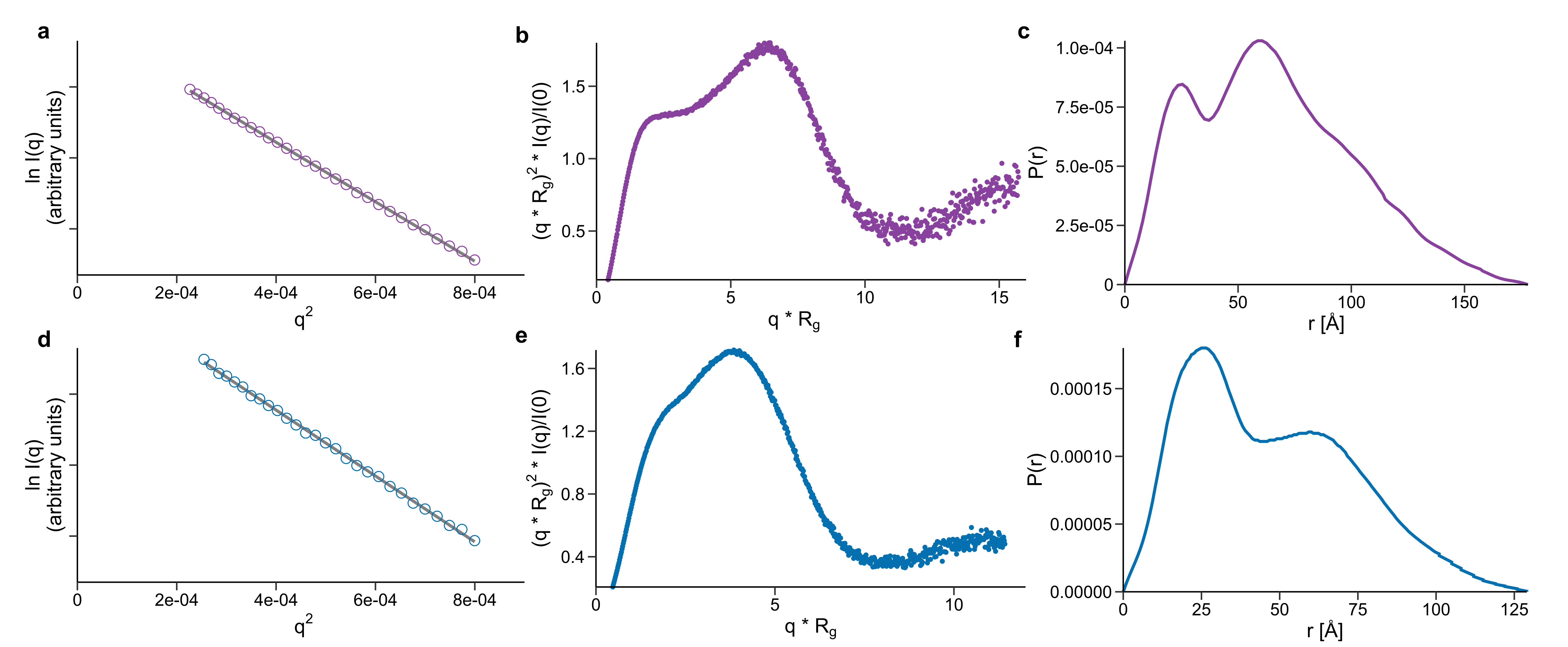

**Fig. S8 SEC-SAXS data.** The experimental SEC-SAXS data were processed and evaluated for the **a-c** Cf-SGBP and the **d-f** Cf-GH16_3. **a+d** Linear Guinier region of the experimental data. The gray line represents the Guinier approximation. **b+c** Normalized Kratky representation. **d+f** Distance distribution function, *p(r)*.

**Table S2.** Details on SEC-SAXS samples, data-collection, analysis and 3D modelling details for Cf-SGBP and Cf-GH16_3

(a) Sample details.	**Cf-SGBP**	**Cf-GH16_3**

Origin	*Christiangramia forsetii*	*Christiangramia forsetii*

GFO_RS17395	GFO_RS16360

M_w_ from chem. comp. (aa)	95559 Da (896aa)	57802 Da (538aa)

Sample environment	10 mM MOPS pH 7.8	10 mM MOPS pH 7.8

100 mM NaCl	100 mM NaCl

Sample temperature (°C)	15	15

In-beam sample cell, flow	1 mm quartz capillary, 0.3 ml/min	1 mm quartz capillary, 0.3 ml/min

Sample concentration(s) (mg/ml)	8.2	12.0

(b) SAXS data collection.

Data-acquisition/reduction software: FOXTROT 3.5.10-3979

Source/instrument description: SOLEIL SWING beamline

Measured q-range (qmin–qmax) (Å^-1^): 0.0036–0.553

Method for scaling intensities: Absolute scaling (cm^-1^) referenced to water

Exposure time(s), No. of exposures 990ms x 880 frames (sample); 990ms x 180 frames (buffer)

(c) SAXS-derived structural parameters.

Method(s)/software: PRIMUS, AUTORG and GNOM (ATSAS 3.1.3 (2)).

Guinier analysis	**Cf-SGBP**	**Cf-GH16_3**

I(0) (cm^-1^)	0.11 ± 0.002†	0.13 ± 0.002†

Rg (Å)	52.34 ± 0.1	38.04 ± 0.7

qRg range (datapoint range)	0.64–1.21 (19–49)	0.56–1.03 (26–53)

Linear fit assessment (AUTORG fidelity)	0.74	0.68

PDDF/P(r) analysis

I(0) (cm^-1^)	0.114 ± 0.001	0.132 ± 0.09

Rg (Å)	52.55 ± 0.11	38.22 ± 0.13

Dmax (Å)	178	129.5

q-range (Å^-1^)	0.012–0.504	0.017–0.504

P(r) reciprocal-space fit/CorMap P-value	1.12, 0.74	1.44, 0.71

(d) Scattering particle size.

Method(s)/software PRIMUS (ATSAS 3.1.3 (2); equation 1 in (3) for M from I(0)/c)

**Cf-SGBP	Cf-GH16_3**

Volume estimates (Å^3^)	104621	67596

Porod volume VP (ratio to M)	96495 (1.0)	55070 (0.95)

M from I(0)/c	86316 (0.9)	55769 (0.97)

(e) Modelling.

Methods/software Dummy-atom (GASBORi), rigid-body modelling with DADIMODO (4) and the Ensemble Optimization Method (EOM) (5)

Shape modelling/software	GASBORi	GASBORi

q-range for fit (Å^-1^)	0.012–0.50	0.0126–0.50

Symmetry/anisotropy assumptions	P1	P1

Normalized spatial discrepancy	0.74	0.86

No. of individual model reconstructions 8 (normalized spatial discrepancy among models 0.74 and 0.86) yielding variable models, but all displayed a bent shape.

(f) Data and model deposition.	**Cf-SGBP	Cf-GH16_3**

SASBDB ID	SASDxxx1	SASDyyy2

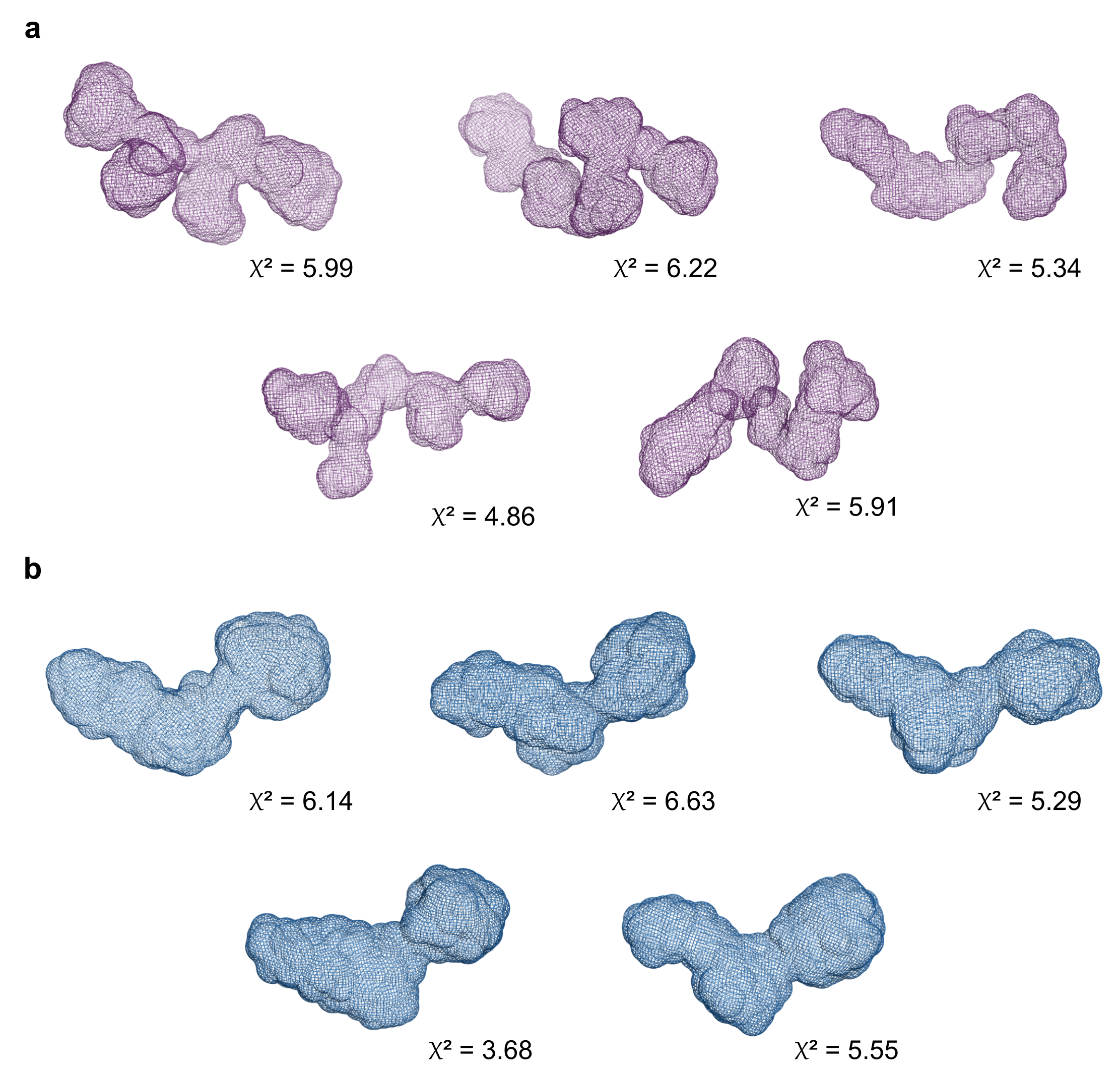

**Fig. S9 Protein envelopes.** GASBORi *ab initio* fitted envelopes for the **a** Cf-SGBP and the **b** Cf-GH16_3. Shown are five calculated models with corresponding χ^2^ values.

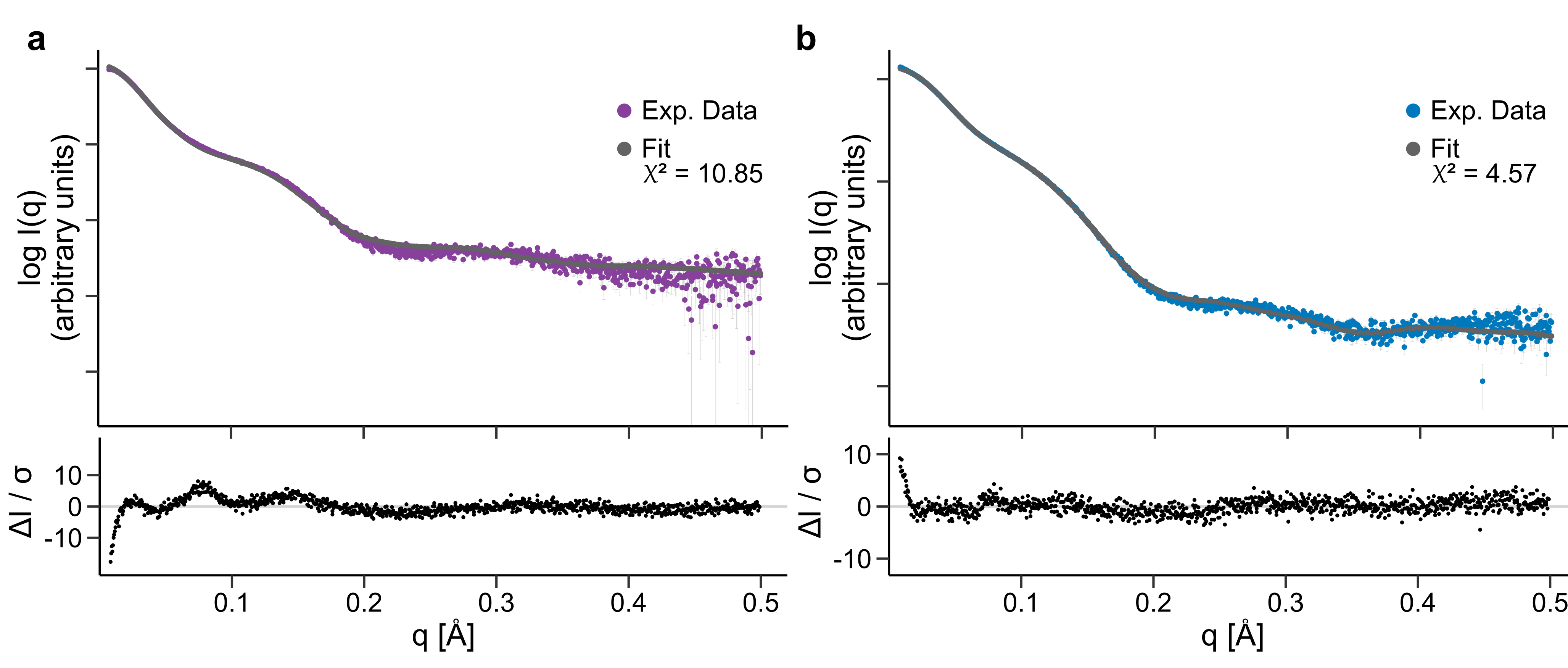

**Fig. S10 Analyzing the flexibility of both multidomain proteins.** EOM was used to investigate the potential conformational flexibility of the **b** Cf-SGBP and the **c** Cf-GH16_3 in solution. Shown are the theoretical scattering curves from the selected ensembles (Fit) plotted against the experimental data (Exp. data).

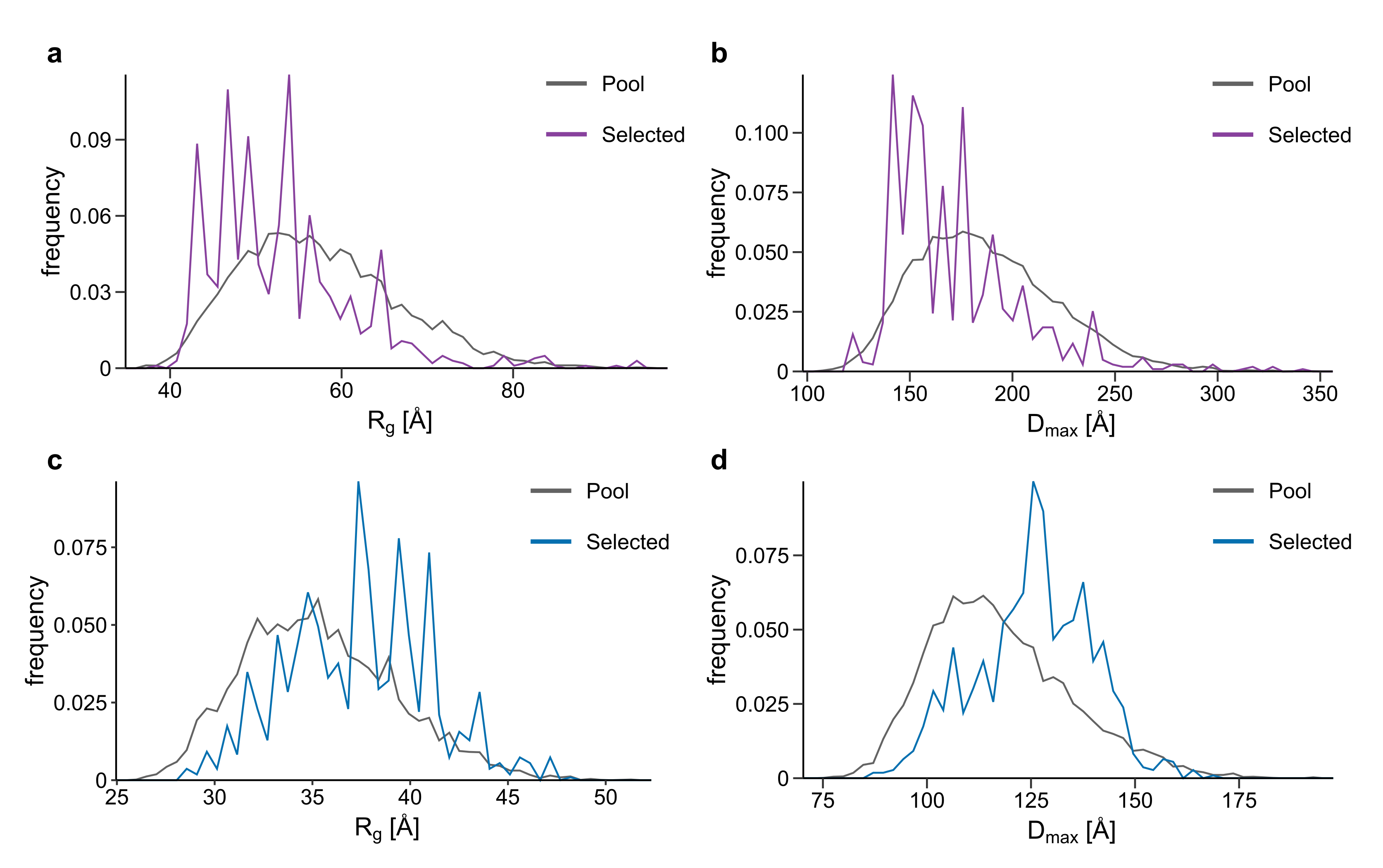

**Fig. S11. Distribution of *R*_g_ and *D*_max_ as determined by EOM analyses.** *R*_g_ and *D*_max_ distributions of the pool (gray line) and selected ensembles (colored line) for the **a+b** Cf-SGBP and the **c+d** Cf-GH16_3. Whereas *R*_g_ and *D*_max_ of selected ensembles were shifted to lower values compared to those of the pool for the Cf-SGBP, they were biased to higher values in the case of the Cf-GH16_3.

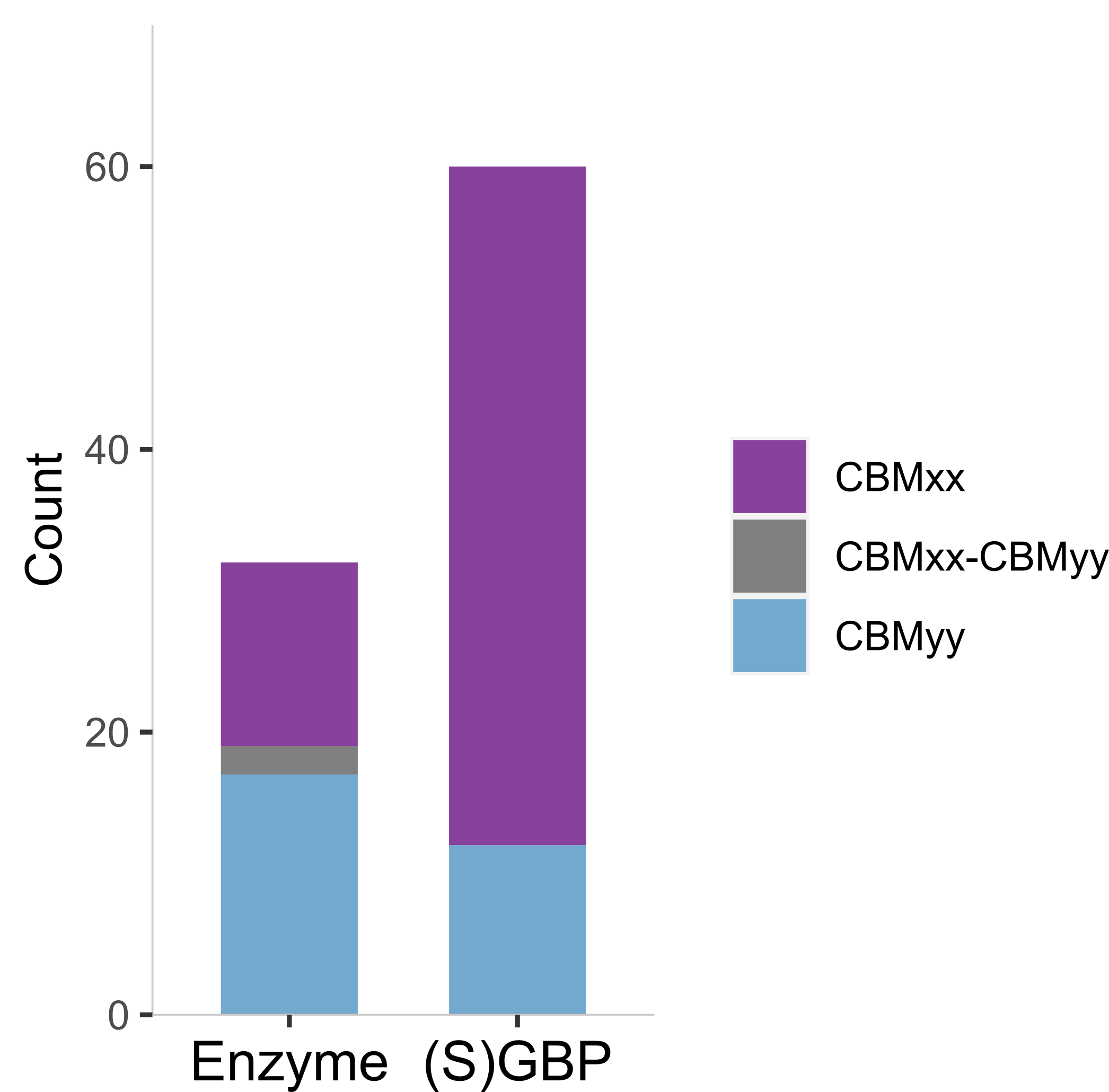

**Fig. S12**. **Distribution of CBMxx- and CBMyy-containing proteins encoded by *Bacteroidota* isolated from the North Sea.** 53 isolates (6) were screened for CBMxx and CBMyy. Shown here are sequences in which at least one CBMxx and/or CBMyy was detected. “Enzyme” refers to their association with GH modules (mostly GH16_3) in polypeptide chains and “(S)GBP” to putative CBM-only proteins.

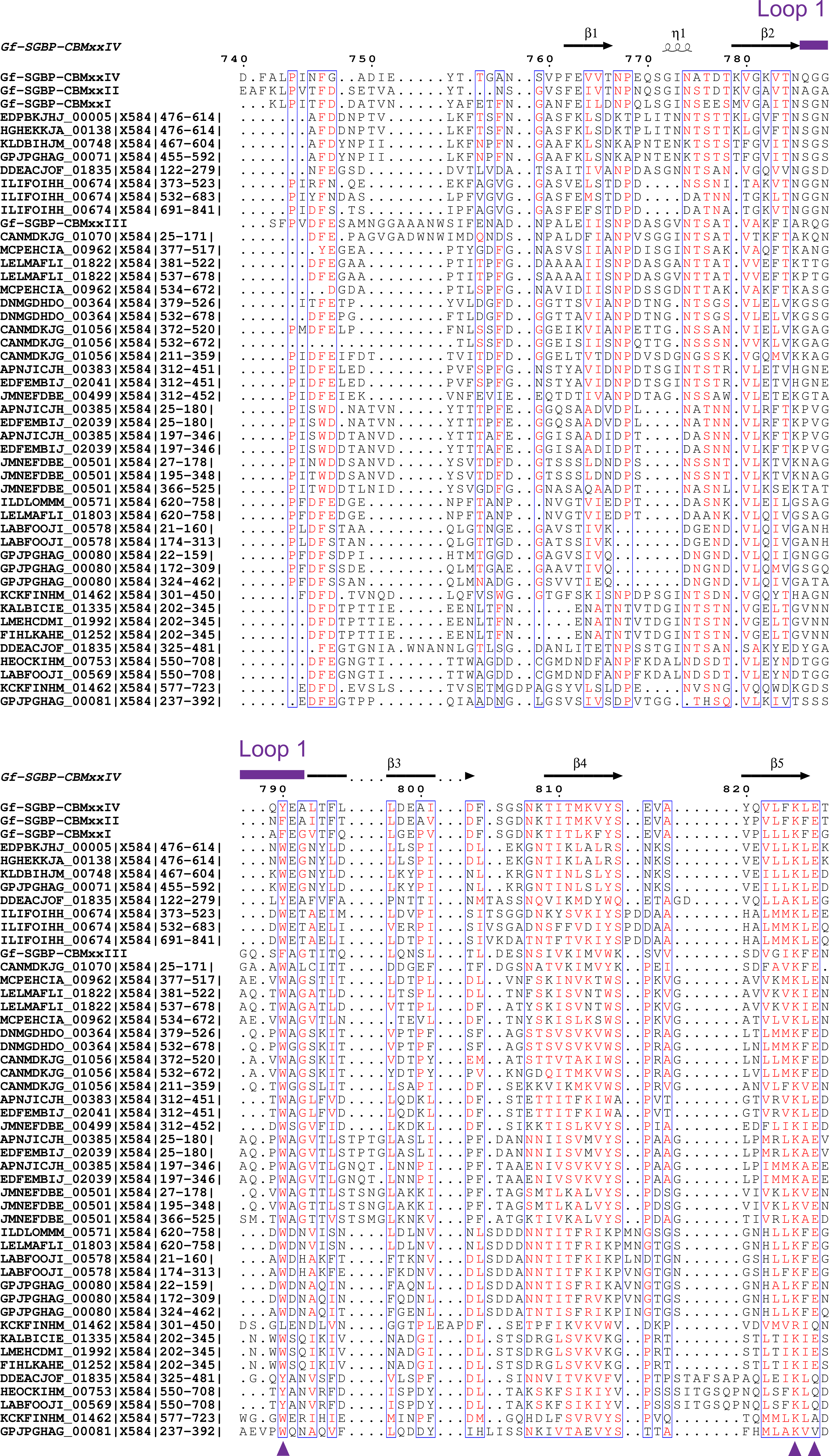

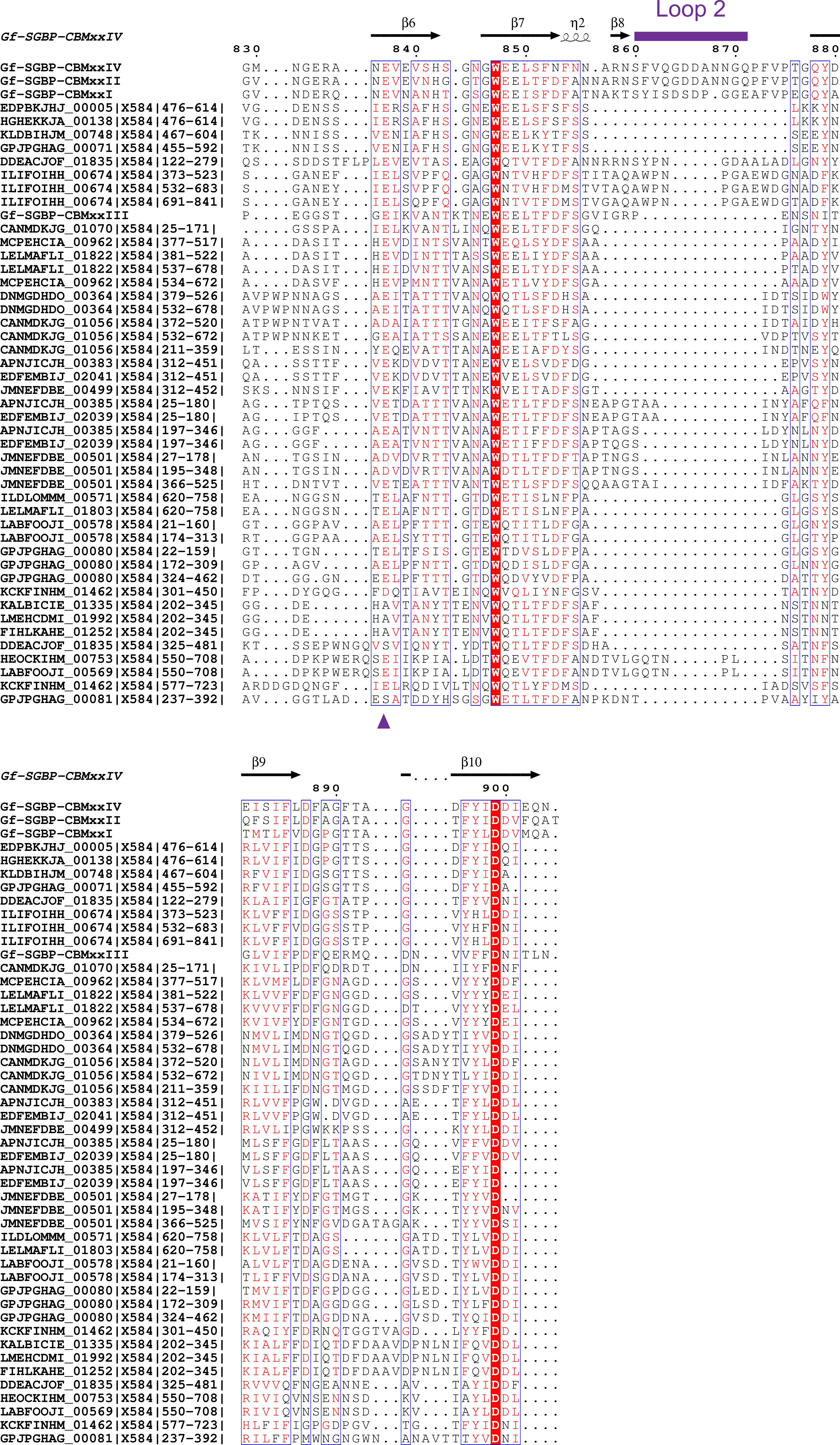

**Fig. S13. Multi-alignment of CBMxxs of the Cf-SGBP and selected CBMxx sequences from environmental MAGs.** Sequences were aligned using MAFFT v6.864 with the L-INS-i strategy (<https://www.genome.jp/tools-bin/mafft>) and visualized using ESPript 3.0 (<https://espript.ibcp.fr>) (7). Residues involved in substrate binding in the case of the Cf-SGBP-CBMxx_IV_ are indicated by violet arrow heads; the two loops flanking the substrate binding groove are indicated by violet lines above the alignment.

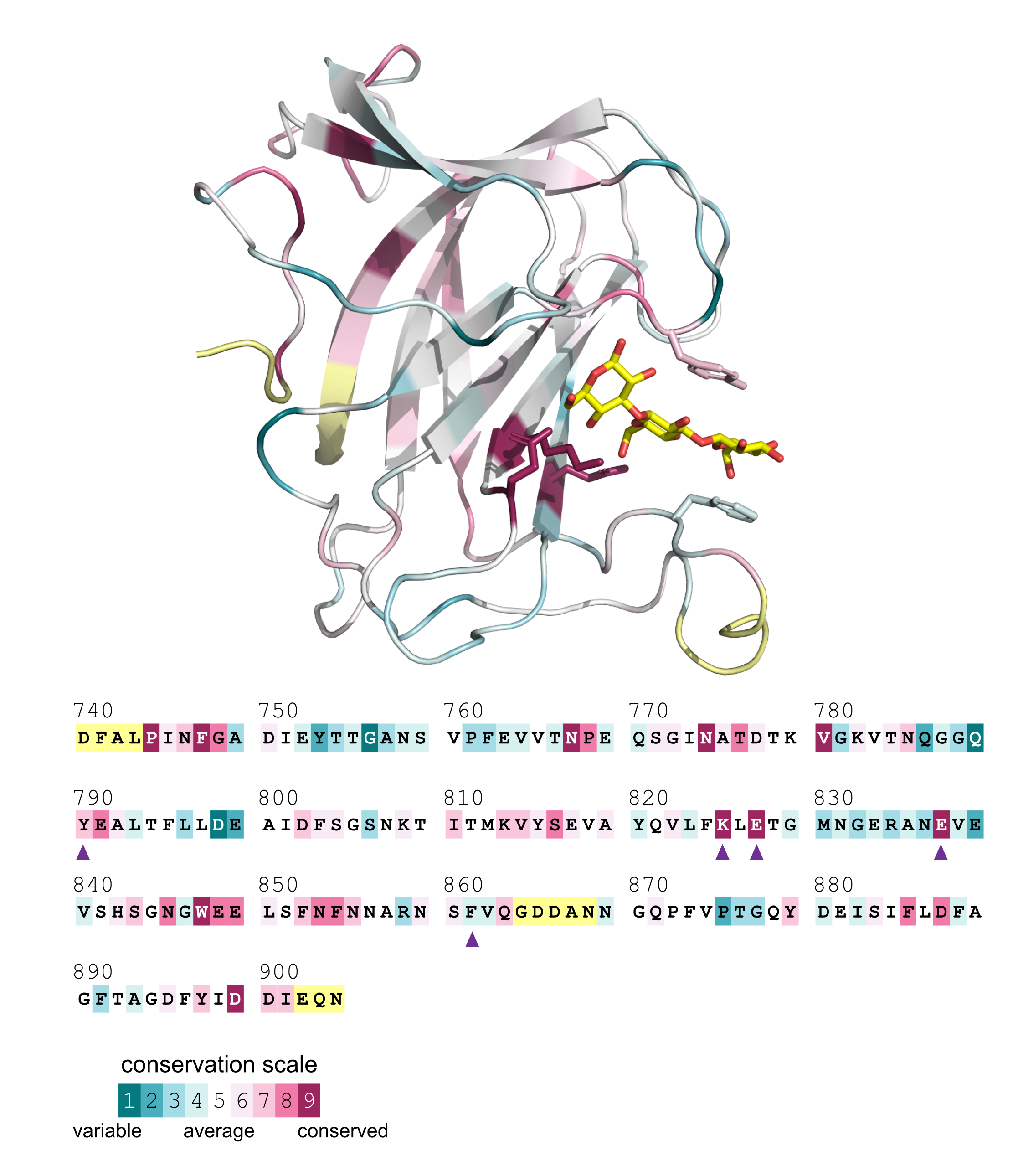

**Fig. S14 Conserved residues of CBMxx.** Conservation of CBMxxs of the Cf-SGBP and selected CBMxx sequences from environmental MAGs was investigated using the ConSurf Server (<https://consurf.tau.ac.il/consurf_index.php>). The analysis was performed using a MAFFT v6.864 alignment with the L-INS-i strategy (<https://www.genome.jp/tools-bin/mafft>), see Fig. S13, and the 3D crystal structure of Cf-SGBP-CBMxx_IV_.

**Table S3 Primer list.** List of primers used for gene amplification and site-directed mutagenesis. Cf-GH16_3-Cat_E442S_ and Cf-GH16_3-Cat_E447S_ are mutants of Cf-GH16_3-Cat, corresponding primers were used to insert the mutation (for details please refer to the Materials and Methods). Nucleotide triplets that replaced the glutamic acids by serines are highlighted in bold and italics. The vector primers anneal to pRF3 and pET28a directly upstream and downstream of the multiple cloning site. Cf-SGBP-Ig_a/b_, Cf-SGBP-CBMxx_I_, Cf-GH16_3-Ig and Cf-GH16_3-CBMyy were ordered.

| Product | Primer | Sequence |
| --- | --- | --- |
| Cf-SGBP | fw | AAAAAAGGATCCgaaaaagaggaagaaacatcatatatac |
| Cf-SGBP | rev | AAAAAACCTAGGCTAATTCTGTTCTATATCATCAATGTAG |
| Cf-GH16_3 | fw | AAAAAAGGATCCgtagacgatgattatgatattggaac |
| Cf-GH16_3 | rev | AAAAAACCTAGGTTATTGATACACTTTGATGTAGTCAATT |
| Cf-SGBP-Ig_a_ | fw | GCGCCATATGGAAAAAGAGGAAGAAACATCATATATACTGC |
| Cf-SGBP-Ig_a_ | rev | CTGTGAGCTCTCAAGATTTTTCTATATCTACCTCAAGTTCTGAC |
| Cf-SGBP-truncated | fw | GCGCCATATGGAAAAAGAGGAAGAAACATCATATATACTGC |
| Cf-SGBP-truncated | rev | CTGTGAGCTCTCATCCGGCAGCATTTAAAGTAATATTATCAAAAAAGAC |
| Cf-SGBP-CBM_II/III_ | fw | CTGTCATATGGGCGAACCTGGAGAAGCATTTAAATTAC |
| Cf-SGBP-CBM_II/III_ | rev | CTGTGAGCTCTCATCCGGCAGCATTTAAAGTAATATTATCAAAAAAGAC |
| Cf-SGBP-CBM_III/IV_ | fw | CTGTCATATGGGTGCCACAGGAATTACTACCACAAGTTTCCCG |
| Cf-SGBP-CBM_III/IV_ | rev | CTGTGAGCTCCTAATTCTGTTCTATATCATCAATGTAGAAATCAC |
| Cf-SGBP-CBM_IV_ | fw | CTGTCATATGGGTGAAACTTCGGAAGATTTTGCGCTGCC |
| Cf-SGBP-CBM_IV_ | rev | CTGTGAGCTCCTAATTCTGTTCTATATCATCAATGTAGAAATCAC |
| Cf-GH16_3-Cat | fw | GCGCCATATGGGAGAAGAAGAATTTGAGACTCAATTTGAAGAC |
| Cf-GH16_3-Cat | rev | CTGTGAGCTCTTATTGATACACTTTGATGTAGTCAATTTCCATAG |
| Cf-GH16_3-Cat_E442S_ | fw | GGATGTTAGGCGCTAATTTTGACGAAGTGGGTTGGCCTGAAACCGGT***TCG***ATCGATATTATG |
| Cf-GH16_3-Cat_E442S_ | rev | GATGAAGTCCTATTTGGCTCATTTCCTACCCATTCCATAATATCGAT***CGA***ACCGGTTTC |
| Cf-GH16_3-Cat_E447S_ | fw | GACGAAGTGGGTTGGCCTGAAACCGGTGAGATCGATATTATG***TCA***TGGGTAGGAAATG |
| Cf-GH16_3-Cat_E447S_ | rev | CAAATGAAGTGCAGATGAAGTCCTATTTGGCTCATTTCCTACCCA***TGA***CATAATATCGATC |
| vector | fw | TAGGGGAATTGTGAGCGG |
| vector | rev | CAAGACCCGTTTAGAGGC |
